# Modulating the voltage sensor of a cardiac potassium channel shows antiarrhythmic effects

**DOI:** 10.1101/2021.02.25.432939

**Authors:** Yangyang Lin, Sam Z. Grinter, Zhongju Lu, Xianjin Xu, Hong Zhan Wang, Hongwu Liang, Panpan Hou, Junyuan Gao, Chris Clausen, Jingyi Shi, Wenshan Zhao, Zhiwei Ma, Yongfeng Liu, Kelli McFarland White, Lu Zhao, Po Wei Kang, Guohui Zhang, Ira S. Cohen, Xiaoqin Zou, Jianmin Cui

## Abstract

Cardiac arrhythmias are the most common cause of sudden cardiac death worldwide. Lengthening the ventricular action potential duration (APD) either congenitally or via pathologic or pharmacologic means, predisposes to a life-threatening ventricular arrhythmia, Torsade de Pointes. IKs, a slowly activating K^+^ current plays a role in action potential repolarization. In this study, we screened a chemical library *in silico* by docking compounds to the voltage sensing domain (VSD) of the I_Ks_ channel. Here we show that C28 specifically shifted I_Ks_ VSD activation in ventricle to more negative voltages and reversed drug-induced lengthening of APD. At the same dosage, C28 did not cause significant changes of the normal APD in either ventricle or atrium. This study provides evidence in support of a computational prediction of I_Ks_ VSD activation as a potential therapeutic approach for all forms of APD prolongation. This outcome could expand the therapeutic efficacy of a myriad of currently approved drugs that may trigger arrhythmias.

**Significance statement:** C28, identified by *in silico* screening, specifically facilitated voltage dependent activation of a cardiac potassium ion channel, I_Ks_. C28 reversed drug-induced prolongation of action potentials, but minimally affected the normal action potential at the same dosage. This outcome supports a computational prediction of modulating I_Ks_ activation as a potential therapy for all forms of action potential prolongation, and could expand therapeutic efficacy of many currently approved drugs that may trigger arrhythmias.

## Introduction

I_Ks_, a slowly activating delayer rectifier in the heart is important in controlling cardiac action potential duration (APD) and adaptation of heart rate in various physiological conditions(1). The I_Ks_ potassium channel has slow activation kinetics and the activation terminates cardiac action potentials (2). This channel is formed by the voltage gated potassium (Kv) channel subunit KCNQ1 and the regulatory subunit KCNE1. The association of KCNE1 drastically alters the phenotype of the channel, including a shift of voltage dependence of activation to more positive voltages, a slower activation time course, a changed ion selectivity, and different responses to drugs and modulators(3–6). Similar to other K_V_ channels, KCNQ1 has six transmembrane segments S1-S6, in which S1-S4 form the voltage sensing domain (VSD), while S5 and S6 form the pore domain; four KCNQ1 subunits comprise the KCNQ1 channel(7, 8). KCNQ1 and I_Ks_ channels are activated by voltage. The VSD in response to membrane depolarization changes conformation, triggered by the movements of the S4 segment that contains positively charged residues (9–15). This conformational change alters the interactions between the vSD and the pore, known as the vSD-pore coupling, to induce pore opening (12–15).

The ventricular APD depends on the balance of outward and inward currents flowing at plateau potentials. The outward currents include the delayed rectifiers I_Kr_ and I_Ks_, while the inward currents include a persistent sodium current (I_NaP_)(16). Specific mutations in any of these channel proteins that cause a reduction in outward current or increase in inward current are associated with congenital long QT syndrome (LQTS), a condition in which the APD is abnormally prolonged, predisposing the afflicted patients to a lethal cardiac arrhythmia called Torsades de Pointes (TdP)(17). In fact, mutations in multiple genes that alter the function of various ion channels have been associated with LQTS(18). There is also a much more prevalent problem called acquired LQTS (aLQTS) that is most often associated with off target effects of drugs. Many drugs are marketed with a QT prolongation warning, and the drug concentrations that can be used therapeutically are limited by this potentially lethal side effect. Some effective drugs have been removed from market(19) because of Q-T prolongation, and others are abandoned before clinical trials even began. Therefore, aLQTS is costly for the pharmaceutical industry both in drug development (to avoid this side effect) and when it results in removal from the market of compounds that have effectively treated other diseases(20, 21). At present, the I_Kr_ (HERG) potassium channel(20) and the phosphoinositide 3-kinase (PI3K)(22, 23) have been identified as the most prominent off targets of these drugs for association with aLQTS.

We hypothesized that in LQT the normal heart function can be restored and Q-T prolongation prevented by compensating for the change in net current from any of the channels produced by the myocyte; all that is required is that a reasonable facsimile of normal net current flow be restored. We applied this approach to aLQTS based on a computational study to show that a shift of voltage dependent activation of I_Ks_ to more negative voltages would increase I_Ks_ during ventricular action potentials; this increase of I_Ks_ would revert drug-induced APD prolongation to normal. More importantly, a change in I_Ks_ voltage dependent activation might affect the normal APD to a much smaller degree due to its slow activation kinetics in healthy ventricular myocytes, thereby posing minimal risk of cardiac toxicity on its own. To apply this approach experimentally, we needed a compound that could specifically shift the voltage dependence of I_Ks_ activation. Previous studies showed that the benzodiazepine R-L3(24) and polyunsaturated fatty acids (PUFAs)(25) activate I_Ks_ channels, while more recently rottlerin was shown to act similarly to R-L3(26). However, the effects of these compounds on I_Ks_ are complex, likely through binding to more than one site in the channel protein instead of simply acting on voltage dependent activation(24, 27, 28). In addition, these compounds showed poor specificity for I_Ks_, also affecting other ion channels in the heart(27, 29, 30), which make these compounds unsuitable to test our hypothesis. Recently, we have identified CP1 as an activator for I_Ks_, which mimics the membrane lipid phosphatidylinositol 4,5-bisphosphate (PIP2) to mediate the VSD-pore coupling(13, 31). CP1 enhances I_Ks_ primarily by increasing current amplitude with some shift of voltage dependence of activation, which is not suitable for our test either.

In this study we identified a compound C28, using an approach that combines *in silico* and experimental screening, that interacts with the KCNQ1 VSD and shifts voltage dependence of vSD activation to more negative voltages. C28 increases both exogenously expressed I_Ks_ and the current in native cardiac myocytes. As predicted by computational modeling, C28 can prevent or reverse the drug induced APD prolongation back to normal, while having a minimal effect on the control APD at the same concentration in healthy cardiac myocytes. This study demonstrates that the KCNQ1 VSD can be used as a drug target for developing a therapy for LQT, and C28 identified in this study may be used as a lead for this development. Further, our results provide support for the use of docking computations based on ion channel structure and cellular physiology, in combination with functional studies based on molecular mechanisms, as an effective approach for rational drug design.

## Results

### C28 shifts voltage dependence of KCNQ1 and I_Ks_ to more negative voltages

In a previous study we showed that, during activation of the KCNQ1 voltage sensing domain (VSD, Fig 1A), residue E160 in the S2 helix interacts with positively charged residues in S4 as they move toward the extracellular side of the membrane in response to depolarization of the membrane potential, and these interactions could be modified by extracellular compounds to alter channel gating(9). This result suggested that the KCNQ1 VSD could serve as a target for developing novel antiarrhythmic drugs. To identify a compound that would bind to the pocket near these interactions, we used our MDock docking software(32–34) to screen *in silico* a compound library targeting the VSD of KCNQ1, as described in the Materials and Methods (see Supplementary Materials). We used the Available Chemical Database (ACD, Molecular Design Ltd.) as the compound library, which consists of over 200,000 organic compounds. The *in silico* screening produced 53 candidates, which were further screened experimentally using voltage clamp. The compound C28 was further docked onto the recently solved cryo-EM structure of human KCNQ1 (8) (Fig 1A, B). C28 was found to cause a shift in voltage dependence of activation for both KCNQ1 and KCNQ1+KCNE1 (termed I_Ks_ thereafter) channels (Fig 1, 2).

**Figure 1.**
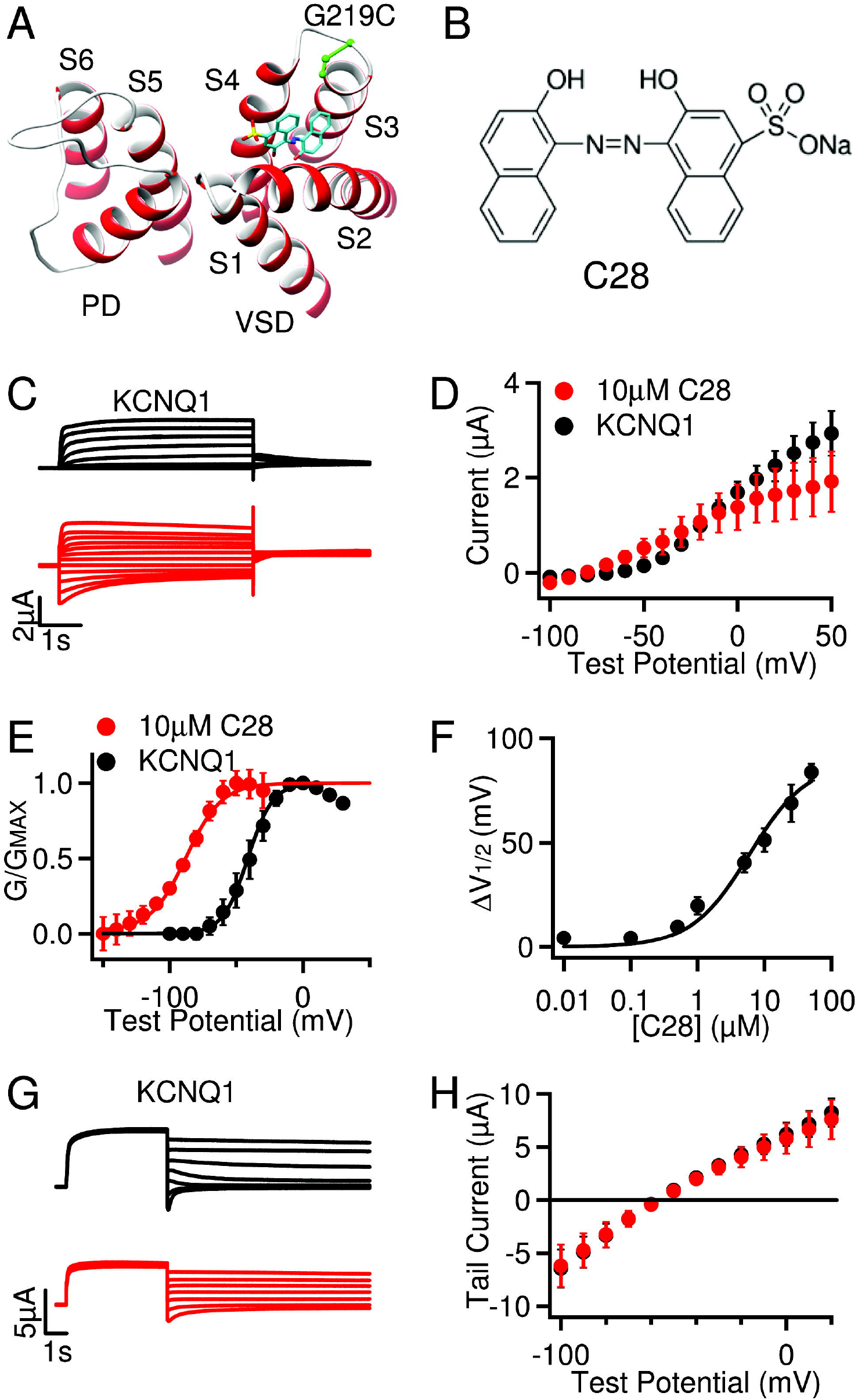
C28 effects on KCNQ1 opening. **A,** KCNQ1 cryo-EM structure (PDB: 6uzz) docking with C28 (cyan, yellow and red sticks). G219C (green) was covalently labeled with Alexa488 in VCF experiments. S1-S6: transmembrane helices. The voltage sensing domain (VSD, S1-S4) and the pore domain (PD, S5-S6) are from neighboring subunits. **B**, C28 molecule. **C,** Currents of KCNQ1 (black) and with C28 (10 μM, red) at various test voltages (see **D**). The potentials before and after test pulses were −80 and −40 mV, respectively. **D**, Steady-state current-voltage relations, with current amplitudes at the end of test pulses shown in C. p>0.05 between control and C28 at all voltages, unpaired student test. **E,** G-V relations. Solid lines are fits to the Boltzmann relation with V1/2 and slope factor (mV) for control: −41.4 ± 1.4 and 9.3 ± 1.3; and for 10 μM C28: −87.7 ± 1.4 and 13.3 ± 1.4. **F**, The change of V1/2 of G-V relations depends on C28 concentration, with EC_50_ of 7.6 μM. **G**, Reversal potential measurements. KCNQ1 channels were activated at 40mV and then currents were tested at −100mV to 20mV. Holding potential: −80 mV. Control: black, with C28 (10 μM): red. **H,** Peak tail currents in relation to the test voltages. The currents reversed at potentials (mV): −58.2± 0.8 for control (black) and – 57.1± 1.1 for 10 μM C28 (red). All data in this figure and subsequent figures are mean ± sem, n = 3-15 unless otherwise specified.

**Figure 2.**
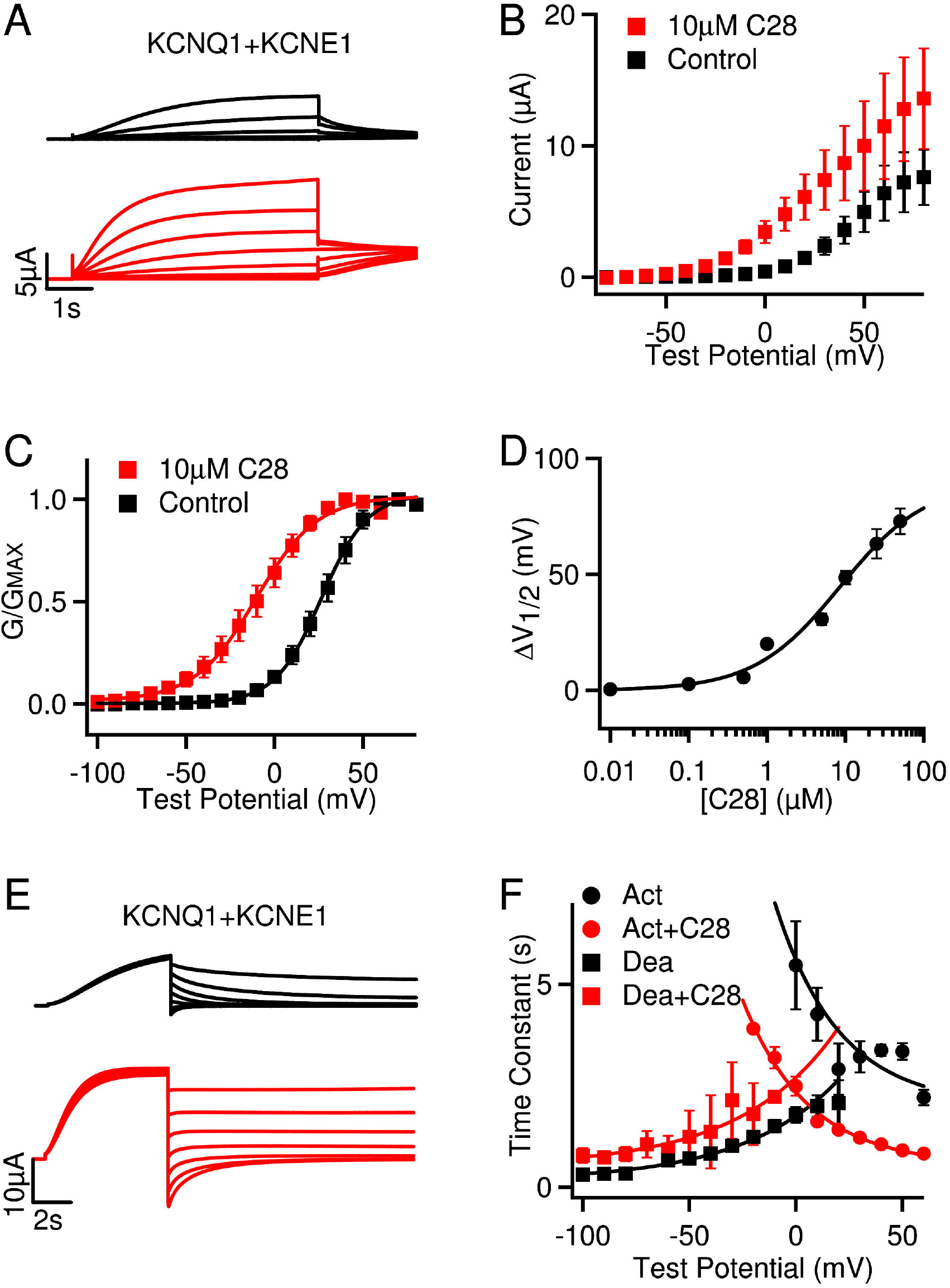
C28 effects on I_Ks_ opening. **A,** I_Ks_ (black) and with C28 (10 μM, red) at various voltages (see **B**). The potentials before and after test pulses were −80 and −40 mV, respectively. **B**, Steady-state current-voltage relations. P<0.05 between control and C28 at voltages −50 – +20 mV, unpaired student test. **C,** G-V relations. Solid lines are fits to the Boltzmann relation with V1/2 and slope factor (mV) 26.4 ± 0.6 and 13.1 ± 0.5 (control) and −10.4 ± 1.3 and 17.3 ± 1.3 (10 μM C28). **D,** G-V shifts in response to C28. EC_50_ is 5.9 μM. **E**, I_Ks_ deactivation. The channels were activated at 60mV and then tested at – 100mV to 40mV. Control: black, with C28 (10 μM): red. **F**, Voltage dependence of activation (Act) and deactivation (Dea) time constants of I_Ks_ in control and 10 μM C28. The time constants were obtained by exponential fitting to current traces. Solid lines are exponential fits to the data. P<0.05 between control and C28 at all voltages, unpaired student test.

We measured the modulation of KCNQ1 channel activation by C28 at various concentrations. The current amplitude was increased at low voltages with the application of C28, but the changes were not significant (Fig 1C, D), while the voltage dependence of activation, measured as the conductance-voltage (G-V) relation, was shifted to more negative voltages (Fig 1E). These results suggest that C28 enhanced channel activation by altering KCNQ1 voltage dependence. The shift of G-V relation could be as large as – 90 mV, and the EC_50_ for the shift was 7.6 μM (Fig 1F). C28 did not alter the reversal potential of KCNQ1 channels, suggesting that K^+^ selectivity of the channel was not affected by C28 (Fig 1G, H). These results are consistent with our docking computations of C28 interacting with the KCNQ1 VSD (Fig 1A).

We then examined the effects of C28 on I_Ks_ (KCNQ1 + KCNE1) channels. We found that the KCNQ1 and I_Ks_ channel inhibitor XE991 could still inhibit the I_Ks_ channel in the presence of C28 (Fig S1A, B), and the result indicates that C28 did not elicit any change in the endogenous currents in oocytes. C28 also increased current amplitude (Fig 2A, B), and the effects could be washed out (Fig S1C). C28 shifted the G-V relation of the I_Ks_ channel to more negative voltages (Fig 2C), similarly to the result for KCNQ1. At 10 μM C28, the G-V shift was ~ −37 mV for I_Ks_, as compared to ~ −46 mV for KCNQ1 (Fig 1E, 2C). The shift of the I_Ks_ G-V relation showed a similar dependence on C28 concentrations as that of KCNQ1, with the EC_50_ at 5.9 μM (Fig 2D, S1D) as compared to 7.6 μM for KCNQ1 (Fig 1F). Interestingly, the VSD of the KCNQ1 channel activates in two steps, first to an intermediate state and then to the activated state; and the KCNQ1 channel opens predominantly when the VSD is at the intermediate state, while the I_Ks_ channel opens exclusively when the VSD is at the activated state (35, 36). This mechanism is manifested in the result that the G-V relation of KCNQ1 was in a more negative voltage range than the G-V relation of I_Ks_ (Fig 1E, 2C). Therefore, the similar shift of G-V relations for both KCNQ1 and I_Ks_ caused by C28 suggest that C28 alters VSD activation to both intermediate and activated states, which is supported by the direct studies of VSD activation (see below). C28 accelerated activation kinetics of I_Ks_, and reduced the rate of deactivation to a small degree (Fig 2E, F).

### C28 modulates KCNQ1 VSD activation by interacting with the VSD

The effects of C28 on the voltage dependence of KCNQ1 and I_Ks_ opening are consistent with our docking result that C28 may bind to the VSD of the channels and alter VSD activation. We further validated this mechanism by first measuring the effects of C28 on VSD activation and then identifying the residues in the KCNQ1 VSD that may interact with C28. To directly examine if C28 modulates the VSD, we measured VSD activation and channel opening in response to voltage changes using voltage-clamp fluorometry (VCF) (Fig 3). The fluorophore, Alexa 488 C5 maleimide, labeling C219 in the VSD (Fig 1A) of the KCNQ1 pseudoWT (psWT) channels (carrying mutations C214A/G219C/C331A), reported VSD movements, while the ionic current reported pore opening(10–14, 36). The mutations in the psWT avoid nonspecific labeling of native Cys214 and Cys331. These mutations and fluorophore labeling had a small effect on KCNQ1 activation, shifting the voltage dependence of channel activation by ~ 9 mV to more negative voltages in the absence of C28 (Fig 1E, 3A, B). However, similar to the WT KCNQ1 channel, C28 also facilitated opening of psWT KCNQ1 channels by shifting the G-V relation to more negative voltages (Fig 3B). C28 caused a striking change of the fluorescence emission, such that instead of an increasing of fluorescence in response to VSD activation, the fluorescence decreased in response to VSD activation upon the wash in of C28 (Fig 3C). The change of fluorescence during each voltage pulse is a result of the environmental change surrounding the fluorophore due to VSD movements in response to voltage(13, 14). The fluorophore was labeled close to the target pocket in our *in silico* screening (C219, Fig 1A). C28 reversed the direction of the fluorescence change during VSD activation (Fig 3C, D), supporting the hypothesis that C28 binds to the targeted pocket in the VSD (Fig 1A) and alters the fluorophore in relation to its environment during VSD movements, by changing either the fluorophore or the conformation of the VSD. In fact, C28 might have affected the fluorophore severely, such that at C28 concentrations higher than 1.5 μM used in these experiments (Fig 3) the VCF recordings were not stable and could not be completed. We measured the steady state voltage dependence of the change of fluorescence emission, *ΔF/F* (Fig 3D, E), which increased with voltage in two distinct steps (Fig 3E), reflecting the step-wise VSD movements of KCNQ1 from the resting state to the intermediate state and then to the activated state(9–12, 35–37). C28 shifted both components of the Δ*F/F*-V relation to more negative voltages (Fig 3E), consistent with similar C28 effects on both KCNQ1 and I_Ks_ opening (Fig 1, 2). These results indicate that C28 facilitates VSD activation.

**Figure 3.**
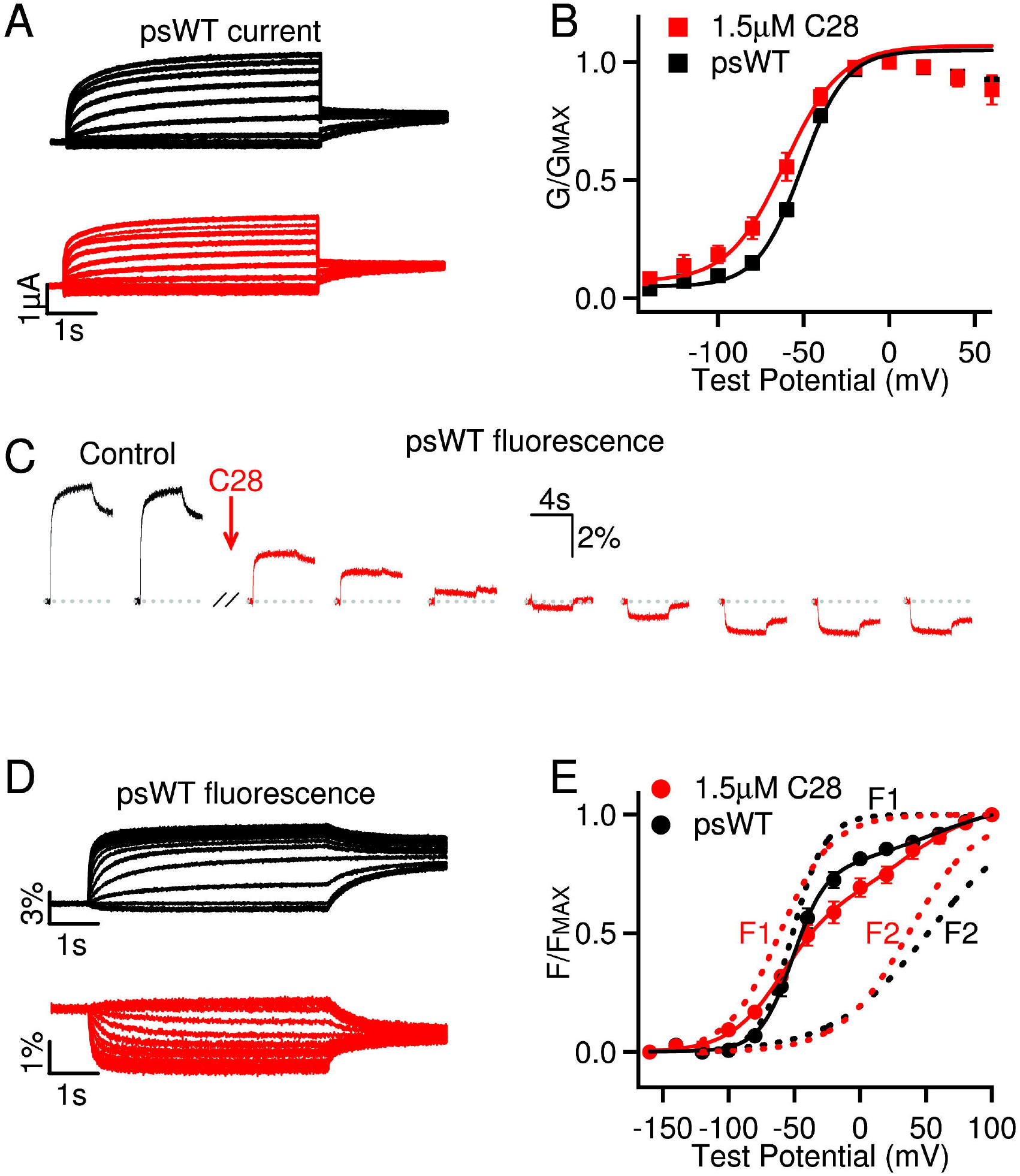
C28 enhances KCNQ1 VSD activation. **A,** Currents of psWT KCNQ1 (black) and with C28 (1.5 μM, red) at various test voltages (see **B**). The potentials before and after test pulses were −80 and −40 mV, respectively. **B**, G-V relations. Solid lines are fits to the Boltzmann relation with V1/2 and slope factor (mV) −50.4 ± 0.9 and 13.9 ± 0.7 (psWT) and −59.7 ± 1.1 and 15.2 ± 1.4 (C28), n = 7. **C**, Fluorescence change of psWT in response to voltage pulses to 40 mV (−80 mV before and −40 mV after the test pulse, pulse interval: 40 s) altered direction upon C28 (1.5 μM) application. **D,** Fluorescence changes of psWT KCNQ1 (black) and with C28 (1.5 μM, red) in response to the voltage protocols from the same oocyte in **A**. **E**, Normalized fluorescence-voltage relation. The curves are fits with double Boltzmann functions (F1, F2) with the V1/2 (mV) F1, −51.9 ± 1.1 and F2, 49.1 ± 2.0 (psWT); and F1, −60.6 ± 1.4 and F2, 39.1 ± 3.3 (C28); and the slope factor (mV): F1, 12.2± 0.5 and F2, 36.0 ± 0.8 (psWT); and F1,21.5 ±1.4 and F2 26.5 ±3.1 (C28); n = 5. The statistical significance of differences in V1/2 between C28 and control were tested using unpaired student test, p < 0.001 for G-V (**B**); p < 0.005 and p < 0.05 for F1 and F2, respectively (**E**).

To examine if C28 interacts with the KCNQ1 VSD, we first measured C28 effects on Kir1.1, which is a two-transmembrane-segment (2TM) K^+^ channel that contains a PD but not the VSD. C28 had no effect on Kir1.1 channels (Fig S2A, B), consistent with the hypothesis that C28 only interacts with the VSD. All voltage gated potassium (KV) channels have VSDs that share a similar structure containing four transmembrane helices S1-S4. However, we found that, unlike its action on KCNQ1, C28 did not alter voltage dependent activation of the Shaker K^+^ channel (Fig S2C, D). To test if this result derives from a specificity of C28 to the KCNQ1 VSD, we studied the KTQ and KTV channels, which are comprised of the two-pore-domain channel TASK3 fused with the KCNQ1 and KV1.2 VSD, respectively(38). TASK3 itself lacks a VSD so that its opening is not dependent on voltage(39), but Lan et al.(38) demonstrated that the fused VSD induces voltage dependent gating in the KTQ and KTV channels (Fig 4A-D). We found that C28 shifted voltage dependent activation of the KTQ channel, which is comprised of TASK3 fused with the KCNQ1 VSD, to more negative voltages and suppressed the amplitude of the current (Fig 4A, B). However, C28 had no effect on the KTV channel, which is comprised of TASK3 fused with the Kv1.2 VSD (Fig 4C, D). These results suggest that the VSD of KCNQ1 specifically renders C28 sensitivity to the KTQ channel. The VSDs of Shaker and its ortholog Kv1.2 do not respond to C28.

**Figure 4.**
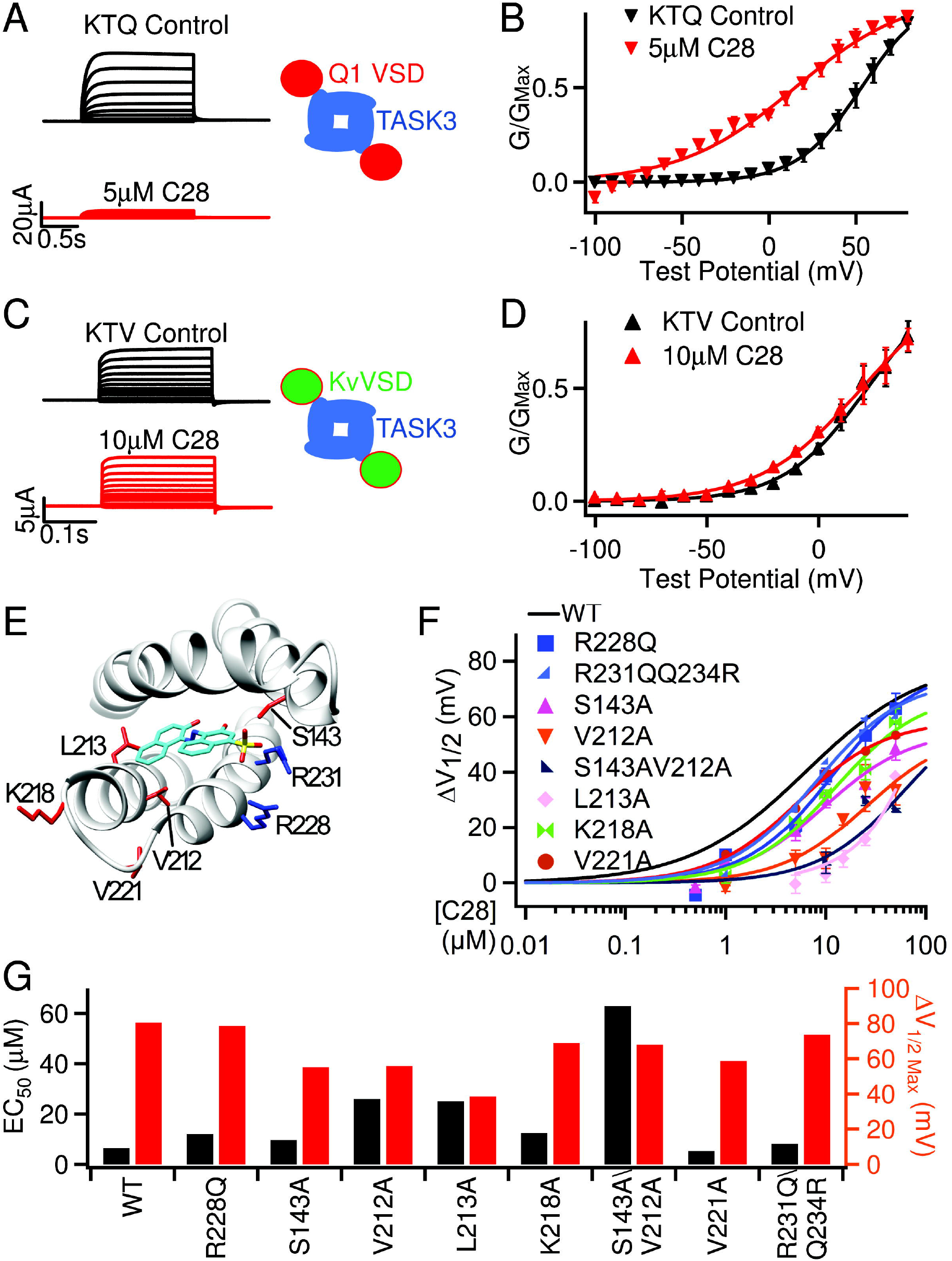
C28 interactions with the VSD of KCNQ1. **A, B**, C28 alters KTQ channel activation. KTQ is a fusion protein between the two-pore domain channel TASK3 and KCNQ1 VSD (inset). Currents were elicited by voltage pulses from −100 to 80 mV with 20 mV increments from a holding potential of −80 mV (**A**). G-V and Boltzmann fits (solid lines) with V1/2 and slope factor (mV) for control: 53.1 ± 2.5 and 18.4 ± 2.5; and for C28: 14.9± 8.2 and 32.8 ± 8.0 (**B**). **C, D**, C28 has no effect on KTV channel activation. KTV is a fusion protein between TASK3 and Kv1.2 VSD (inset). Currents were elicited by voltage pulses from −100 to 40 mV with 20 mV increments(**C**). G-V and Boltzmann fits (solid lines) with V1/2 and slope factor (mV) for control: 21.7 ± 4.0 and 18.5 ± 4.2; and for C28: 19.2± 3.8 and 22.5 ± 4.0 (**D**). **E**, Residues interacting with C28 based on molecular docking to the cryo-EM structure of human KCNQ1 (PDB entry 6uzz). The S3-S4 loop was missing in the cryo-EM structure, and we built it using MODELLER (55). **F,** G-V shift of mutant I_Ks_ in response to C28. Each data point was averaged from recordings in 3-7 oocytes. The curve for WT is the same as in Fig 2D. **G**, Maximal G-V shift and EC_50_ for each mutation and WT.

To further validate the docking results, we identified the residues in the KCNQ1 VSD that are important for C28 modulation by mutagenesis studies. Docking of C28 onto the VSD pocket in the cryo-EM structure of human KCNQ1(8) showed residues that may interact with C28 (Fig 4E, S3A). We mutated these residues to Alanine (except for R228 to Glutamine) and measured the shift of voltage dependent activation of the mutant I_Ks_ channel at various C28 concentrations (Fig 4F). The mutation of these residues increased the C28 concentration for half maximal shift of G-V relations, EC_50_, and decreased the maximum amplitude of G-V shifts (Fig 4G). These include arginine residues in S4 that are part of the voltage sensor, and some of the mutations altered voltage dependent activation of I_Ks_ in the absence of C28 (Fig S3B-D), supporting the idea that C28 binds to our targeted pocket and interferes with voltage sensor movements. A structural comparison between the VSDs of KCNQ1 and K_V_1.2 (Fig S4A, B) shows that the KCNQ1 VSD has short extracellular S1-S2 and S3-S4 loops, which expose the binding pocket completely to the extracellular solution. In addition, the residues that are important for C28 interaction in the KCNQ1 VSD are not conserved in the K_V_1.2 VSD (Fig S4B). These structural differences may underscore the specificity of C28 for KCNQ1 and I_Ks_.

### C28 enhances VSD-pore coupling in I_Ks_ channels

The above results show that C28 activated KCNQ1 and I_Ks_ channels by shifting voltage dependent opening to more negative voltages (Fig 1,2) due to the enhancement of VSD activation by C28 (Fig 3) via interactions with the VSD (Fig 4). However, C28 may activate I_Ks_ via an additional mechanism. The amplitude of I_Ks_ increased with C28 application at all voltages (Fig 2A, B). Even at very positive voltages where the voltage dependent activation was saturated the I_Ks_ amplitude seemed to increase in the presence of C28 (Fig 2B). This suggests an enhancement of the maximum conductance in addition to the shift of the G-V relation. A change in the maximum conductance could be due to a change in the single channel conductance. Since the total number of channels expressed in a cell is not expected to change with C28 application the maximum conductance is proportional to the single channel conductance. However, given that C28 binds to the VSD of I_Ks_ channels but not in the pore (Fig 4), it is unlikely that C28 directly changed single channel conductance of these channels. On the other hand, a change of the VSD-pore coupling could change the efficiency of pore opening in response to VSD activation to alter the maximum conductance. C28 may enhance the efficiency of VSD-pore coupling such that the maximal open probability of each channel is increased to give rise to our observed results.

On the other hand, C28 did not increase the maximum conductance of KCNQ1 (Fig 1C, D). Then why does C28 show different effects on the maximum currents of KCNQ1 and I_Ks_? It has been shown that the KCNQ1 channel can open when the VSD moves to either the intermediate or the activated states, resulting in two distinct open states, the intermediate-open (IO) and the activated-open (AO). For the KCNQ1 channel IO is the predominant open state, whereas for the I_Ks_ channel AO is the exclusive open state because the association of KCNE1 suppresses IO(35, 36). At the IO and AO states the interactions between the VSD and the pore differ, resulting in different VSD-pore coupling(12). In addition, the VSD-pore coupling in KCNQ1 and I_Ks_ channels are dependent on the membrane lipid phosphatidylinositol 4,5-bisphosphate (PIP2)(13, 31), and KCNE1 association enhances PIP2 binding to the channel(4, 36). These results demonstrate that the VSD-pore coupling mechanism differs in the KCNQ1 and IKs channels. Thus, C28 may show different effects on the maximum currents of KCNQ1 and I_Ks_ based on its differential effects on the IO and AO state. To test this idea, we examined if C28 affected the current amplitude of mutant KCNQ1 channels with the VSD arrested in either the intermediate state or the activated state (Fig 5). The mutant KCNQ1 channels were stabilized in the IO and AO states, respectively, producing constitutive currents (Fig. 5A,B, black) and no longer activated by voltage changes (10, 36, 40), Any C28 effect on the current amplitudes of these mutant channels should therefore be proportional to modulation of VSD-pore coupling. We found that C28 enhanced current amplitude in the channel with the VSD arrested in the activated state (Fig 5A, B) but not in the channel with the VSD arrested in the intermediate state (Fig 5C, D). These findings clearly show that C28 modulated VSD-pore coupling of the IO state and AO state differently. Since KCNQ1 and I_Ks_ conduct primarily at the IO and AO state, respectively, these results are consistent with the idea that in the I_Ks_ channel C28 binding to the VSD also affects the interactions between the VSD and the pore, resulting in changes of the VSD-pore coupling to alter the maximum conductance in I_Ks_, but not in KCNQ1.

**Figure 5.**
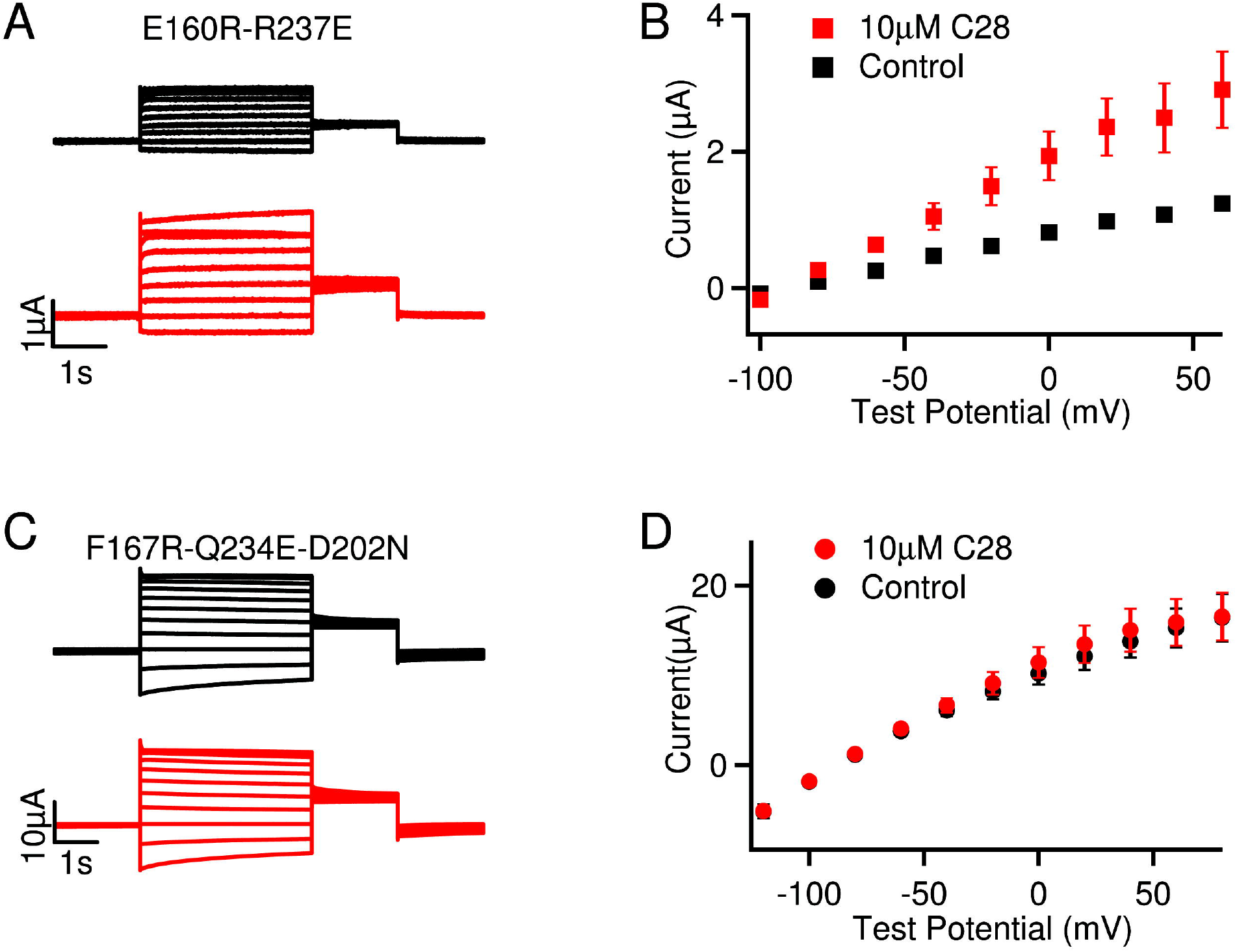
C28 effects on KCNQ1 channels in the activated open (AO) and intermediate open (IO) states. **A,** Current traces of KCNQ1 with mutations E160R- R237E, which arrest the VSD in the activated state and stabilized the channel at the activated open (AO) state (36). Currents were recorded without (black) and with (red) 10 μM C28 at various voltages (see **B**). The voltage before and after the test pulses were – 80 and −40 mV, respectively. **B**, Steady-state current-voltage relations of KCNQ1 E160R- R237E mutant. **C,** Current traces of KCNQ1 with mutations F167R-Q234E-D202N, which arrested the VSD at the intermediate state, thus the channels were stabilized at the intermediate open (IO) state (10). Currents were recorded without (black) and with (red) 10 μM C28 at various voltages (see **D**). The voltage before and after the test pulses were −80 and −40 mV, respectively. **D**, Current-voltage relations of KCNQ1 F167R-Q234E- D202N mutant. P<0.05 in B, but p>0.05 in D between control and C28 at all voltages, unpaired student test.

### C28 reverts drug-induced APD prolongation

Current antiarrhythmic drug therapy restores normal cardiac rhythm by inhibiting membrane ion conductance, which in turn alters the conduction velocity, varies the excitability of cardiac cells by changing the duration of the effective refractory period, and/or suppresses abnormal automaticity(41,42). In a computational simulation of canine ventricular action potentials, we found that an increase of the I_Ks_ current by shifting its voltage dependence of activation to more negative voltages could reverse much of the prolongation of APD caused by the elimination of I_Kr_ (Fig S5A), while inducing only a small shortening in control conditions (Fig S5B). Similar effects of I_Ks_ current on APDs could be observed experimentally on isolated myocytes from canine ventricle in the presence of dofetilide alone, in combination with C28 and with application of C28 alone (Fig S5C, D).

With these encouraging preliminary studies, we further examined the effects of C28 on I_Ks_ and action potentials in cardiac myocytes. We first tested the effects of C28 on other ion channels that are also critical for shaping cardiac action potentials(16) (Fig S6). C28 did not show effects on the voltage dependence of voltage gated Ca^2+^, Na^+^ or other K^+^ channels (Fig S6A-D). These results suggest that, although these channels have VSDs with a similar membrane topology to the KCNQ1 VSD, the specific structural features of the VSD in KCNQ1 and I_Ks_ channels may render it the specificity to its C28 interaction (Fig S4B). On the other hand, C28 activated the M-channel formed by KCNQ2/KCNQ3 with a smaller effect than KCNQ1 (Fig S7), which are expressed in neurons but share a more closely homologous VSD with KCNQ1 (43).

We then tested the effects of C28 on I_Ks_ and action potentials of guinea pig (GP) ventricular myocytes. We used low concentrations of C28 in order to determine the concentration range that might be effective in reducing drug-induced prolongation of action potential duration but less effective in reducing the normal action potential durations. Varying concentrations of C28 (1nM to 1μM) shifted voltage dependence of the I_Ks_ channels in GP ventricular myocytes in a manner similar to those expressed in *Xenopus* oocytes, but in the GP myocytes I_Ks_ seemed to be more sensitive to C28 (Fig 2D, 6A, B). This difference in drug action between oocytes and mammalian cells has been reported for other channels (44). C28 only modulated the currents that were sensitive to the I_Ks_ channel inhibitor Chromanol 293B, but had no effect on currents that were not inhibited by Chromanol 293B (Fig S8A-C), indicating that C28 specifically modified I_Ks_ channels in ventricular myocytes. Furthermore, in ventricular myocytes C28 at these concentrations did not alter the maximum conductance of I_Ks_ in GP ventricular myocytes, but increased I_Ks_ amplitude in the voltage range of the action potential primarily by shifting G-V to more negative voltages (Fig 6A-C, S8D). C28 effectively reversed the prolongation of the APD caused by the I_Kr_ blocker (Moxifloxacin, M), and by a phosphatidylinositol 3- kinase (PI3K) inhibitor (PI-103) (Fig 6D, E). It has been shown that PI-103 decreases I_Kr_ and I_Ks_ currents, the L-type Ca^2+^ channel current I_Ca,L_, and the peak Na^+^ current I_Na_, while it increases the persistent Na^+^ current INaP (22). These results indicate that the enhancement of I_Ks_ currents by C28 stabilized action potentials with alteration of multiple other ion channels. On the other hand, the same dosage of C28 had smaller or absent effects on the normal APD (Fig 6E, S8E, F). Prolonged ventricular action potentials manifest in the whole heart as an increased QT interval in ECG (45). The effects of C28 on APD suggest that C28 should be effective at reducing or eliminating the drug-induced QT prolongation by shifting voltage dependence of I_Ks_ channel activation while having little or no effect in control conditions.

**Figure 6.**
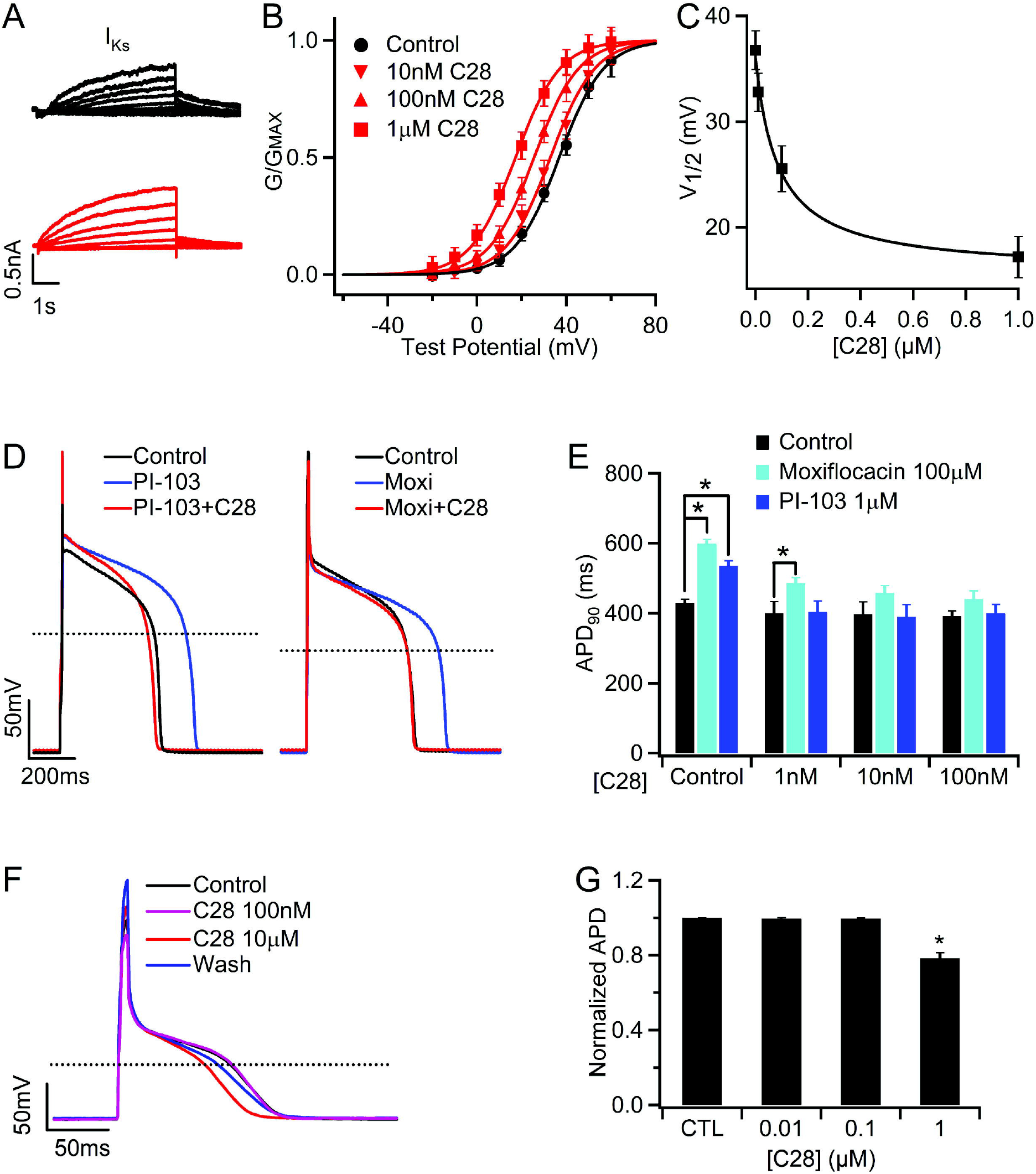
C28 enhances I_Ks_ and stabilizes action potentials in GP ventricular and atrial myocytes. **A,** I_Ks_ currents, measured as the Chromanol 293B sensitive currents (Fig S8A, B), of control (black) and with C28 (100 nM, red) at various voltages (see **B**) from a ventricular myocyte. The voltage before and after the test pulses were −40 and – 20 mV, respectively. **B**, I_Ks_ G-V relations in various C28 concentrations. The lines are fits to Boltzmann equation. **C**, V1/2 of G-V relations versus C28 concentration. **D**, C28 (100 nM) on action potentials in control and the presence of PI-103 (1 μM) and Moxifloxacin (Moxi, 100 μM). **E**, C28 effects on action potential duration (APD). * *P* < 0.05, n = 5-44. **F**, APs of a GP atrial myocyte recorded in control, C28 (100 nM and 10 μM), and after washout (Wash). The stimulus was 180 pA in amplitude, 10 ms in duration at 1Hz frequency. **G**, C28 effects on atrial APD. * P<0.05 compared to control, n = 9-12.

It was shown that some congenital gain-of-function KCNQ1 mutations are associated with atrial fibrillation(46). While C28’s enhancement of I_Ks_ is responsible for reducing drug-induced ventricular APD prolongation, is it prone to induce atrial fibrillation as well? To answer this question, we recorded action potentials in GP atrial myocytes in the presence of various concentrations of C28. The action potential was not affected by C28 up to 1 μM (Fig 6F, G), a concentration that is more than 100 fold higher than the concentration that was effective in reversing drug induced prolongation of ventricular APD (Fig 6D). This result indicates that C28 at a wide range of concentrations may alleviate LQTS symptoms without the risk of causing atrial fibrillation.

## Discussion

The antiarrhythmic drugs currently in clinical use target various ion channels or ß- adrenergic receptors (41,47, 48), but curiously none of these enhances I_Ks_, although the I_Ks_ channel is important in controlling the APD in the heart. In the physiological response to fight or flight heart rate accelerates and I_Ks_ increases to shorten APDs, maintaining stable electrical activity and sufficient diastolic refilling(49). On the other hand, mutations in the I_Ks_ channel subunits, KCNQ1 and KCNE1, predispose patients to multiple types of cardiac arrhythmias(50) such as long QT syndrome(51), short QT syndrome(52), and atrial fibrillation(46). In this study molecular docking studies indicate that a shift of voltage dependent activation of I_Ks_ to more negative voltages is desirable for antiarrhythmic effects with minimal impact on the normal APD (Fig S5). We used *in silico* screening that targeted the VSD of KCNQ1 to identify C28, which was shown experimentally to shift voltage dependence of I_Ks_ activation to more negative voltages and enhance current amplitude (Fig 1–5). C28 is specific in modulating I_Ks_ VSD activation and effective in preventing drug induced APD prolongation or reversing it towards normal in ventricular myocytes (Fig 6). Importantly, C28 shows smaller or negligible effects on APD of normal myocytes from both ventricle and atrium (Fig 6).

For our *in silico* screening we used the KCNQ1 VSD as a target in the hope that the compound docking to the site would directly affect VSD activation. Several lines of evidence validated that C28 is such a compound. We found that C28 shifts voltage dependence of KCNQ1 and I_Ks_ opening (Fig 1,2). C28 interferes with the interactions of gating charge with local environment during charge movements and shifts voltage dependence of VSD activation to more negative voltages (Fig 3). C28 modulation is portable with the KCNQ1 VSD when it is fused with the non-voltage sensitive two-pore-domain channel TASK3 (Fig 4). Mutations of the amino acids that were predicted to interact with C28 in the *in silico* screening reduced C28 modulation (Fig 4). Our study also showed that, although the VSDs in all voltage gated ion channels share a similar structure, C28 is specific for KCNQ1 VSD activation in ventricular myocytes (Fig 4, 6, S2, S6, S8). These results indicate that the KCNQ1 VSD can be used as a drug target to develop antiarrhythmic drugs.

In our study the action potential in GP ventricular myocytes were prolonged by Moxifloxacin and PI-103, which alter multiple cardiac ion channels to cause aLQTS(22). C28 effectively prevented or reversed these effects of the LQT-causing drugs (Fig 6C, D), supporting the hypothesis that increasing I_Ks_ can correct heart rhythm disturbed by changing other ion channels. Drugs approved to treat arrhythmias by blocking specific ion channels lack in efficacy and also have significant proarrhythmic potential(20, 53). For ventricular tachyarrhythmias in particular, antiarrhythmic drug therapy may be contraindicated(54). C28 enhances I_Ks_ to show antiarrhythmic effects, which indicates a novel mechanism on a new target for antiarrhythmic drug development. Importantly, our study shows that C28 has prominent features for cardiac safety. First, C28 affects APD in ventricular myocytes more prominently when it is abnormally prolonged than in control conditions (Fig 6, S5, S8). Second, C28 specifically activates the KCNQ1 VSD and has little effect on the voltage dependence or current of other ion channels that are important in shaping ventricular action potential (Fig 4, S6, S8). The specificity of C28 will exclude off-target effects that may adversely alter heart rhythm. Third, C28 has small effects on atrial APD (Fig 6), reducing the risk of atrial fibrillation as a side effect. These safety features of C28 are encouraging, considering the low concentration required for antiarrhythmic effects, which ranges from 1 to 100 nM in GP ventricular myocytes (Fig 6). At these concentrations, C28 did not alter currents other than I_Ks_, or APD in ventricular or atrial myocytes (Fig 6, S8).

These results describe the development and efficacy of a lead compound for treatment of congenital or acquired long Q-T syndrome (aLQTS). The results are significant, if this approach leads to a drug that can treat congenital and aLQT, for various reasons. 1) The arrhythmia TdP is often lethal. 2) The approach would eliminate an enormous expense in developing new drugs for any therapeutic application because of the FDA requirement that all new drugs minimize risk of aLQT. 3) it would result in greater efficacy for existing drugs, by eliminating current dose restrictions induced by the aLQT side effect. In summary, it would dramatically reduce the cost of drug development, expand current drug efficacy, and save lives.

At present, the ion channels contributing to the function of many specific cells, the diseases associated with ion channel dysfunction in excitable and some non-excitable tissues, the atomistic structures of ion channel families, and the molecular mechanisms of ion channel function are being elucidated at a rapid pace. Meanwhile, more powerful computational hardware and algorithms are also being developed at an increasing speed. This progress may provide a unique opportunity for rationally designed strategies for drug screening and development. This study provides an example to demonstrate the effectiveness of this approach.

## Materials and Methods

C28 was identified by screening a compound library using molecular docking software and the structure model of the KCNQ1 channel, and subsequently testing the effects of the compound on the currents of KCNQ1 and I_Ks_ expressed in *Xenopus* oocytes. In the oocytes, we used the two electrode voltage clamp and voltage clamp fluorometry (VCF) to study the functional effects and molecular mechanism of action of C28 on KCNQ1 and I_Ks_ channels. The effects of C28 on I_Ks_ were examined in freshly isolated Guinea Pig ventricular myocytes while cardiac action potentials were recorded from both Guinea pig ventricular and atrial myocytes using the patch clamp technique.

Detailed methods can be found in Supplementary Materials.

## Acknowledgement

We would like to thank Zhaobin Gao for providing the cDNA of KTQ and KTV fusion proteins. Nien-Du Yang at Washington University helped in some experiments and data analyses. This work was supported by grants R01 HL126774 (JC, XZ and ISC), R01 DK108989 (ISC) and R01 GM109980 (XZ), and R35GM136409 (XZ) from NIH, 13GRNT16990076 (XZ and JS) from AHA (Midwest Affiliate). YL was a visiting student supported by China Scholarship Council (201206380052) via the Sixth Affiliated Hospital of Sun Yat-sen University, China; SZG was also supported through the NLM Biomedical Informatics Research Training Program T15 LM07089. The computations were performed on the high performance computing infrastructure supported by NSF CNS- 1429294 (PI: Chi-Ren Shyu) and the HPC resources supported by the University of Missouri Bioinformatics Consortium (UMBC).

## Author Contributions

Y.L., H.L., P.H., J.S., W.Z., Y.L., K.W., L.Z., P.K., G.Z. and J.C. performed experiments and analyzed data on ion channels expressed in *Xenopus* oocytes. S.Z.G., X.X., Z.M. and X.Z. performed molecular modeling and *in silico* screening. Z.L., H.Z.W., J.G., C.C. and I.S.C. performed experiments and analyzed data on guinea pig and canine ventricular myocytes. J.C., G.Z., I.S.C., and X.Z wrote the manuscript.

## References

1. Silva J & Rudy Y (2005) Subunit interaction determines IKs participation in cardiac repolarization and repolarization reserve. Circulation 112(10):1384–1391.

2. Keating MT & Sanguinetti MC (2001) Molecular and cellular mechanisms of cardiac arrhythmias. Cell 104(4):569–580.

3. Wang HS, Brown BS, McKinnon D, & Cohen IS (2000) Molecular basis for differential sensitivity of KCNQ and I(Ks) channels to the cognitive enhancer XE991. Mol Pharmacol 57(6):1218–1223.

4. Li Y, et al. (2011) KCNE1 enhances phosphatidylinositol 4,5-bisphosphate (PIP2) sensitivity of IKs to modulate channel activity. Proc Natl Acad Sci USA 108(22):9095–9100.

5. Barhanin J, et al. (1996) K(V)LQT1 and lsK (minK) proteins associate to form the I(Ks) cardiac potassium current. Nature 384(6604):78–80.

6. Sanguinetti MC, et al. (1996) Coassembly of K(V)LQT1 and minK (IsK) proteins to form cardiac I(Ks) potassium channel. Nature 384(6604):80–83.

7. Sun J & MacKinnon R (2017) Cryo-EM Structure of a KCNQ1/CaM Complex Reveals Insights into Congenital Long QT Syndrome. Cell 169(6):1042–1050 e1049.

8. Sun J & MacKinnon R (2020) Structural Basis of Human KCNQ1 Modulation and Gating. Cell 180(2):340–347 e349.

9. Wu D, et al. (2010) State-dependent electrostatic interactions of S4 arginines with E1 in S2 during Kv7.1 activation. J Gen Physiol 135(6):595–606.

10. Taylor KC, et al. (2020) Structure and physiological function of the human KCNQ1 channel voltage sensor intermediate state. eLife 9.

11. Barro-Soria R, et al. (2014) KCNE1 divides the voltage sensor movement in KCNQ1/KCNE1 channels into two steps. Nature communications 5:3750.

12. Hou P, et al. (2020) Two-stage electro-mechanical coupling of a KV channel in voltage-dependent activation. Nature communications 11(1):676.

13. Zaydman MA, et al. (2013) Kv7.1 ion channels require a lipid to couple voltage sensing to pore opening. Proc Natl Acad Sci U S A 110(32):13180–13185.

14. Osteen JD, et al. (2010) KCNE1 alters the voltage sensor movements necessary to open the KCNQ1 channel gate. Proc Natl Acad Sci U S A 107(52):22710–22715.

15. Cui J (2016) Voltage-Dependent Gating: Novel Insights from KCNQ1 Channels. Biophys J 110(1):14–25.

16. Nerbonne JM & Kass RS (2005) Molecular Physiology of Cardiac Repolarization. Physiological reviews 85(4):1205–1253.

17. George AL, Jr. (2014) Recent genetic discoveries implicating ion channels in human cardiovascular diseases. Current opinion in pharmacology 15:47–52.

18. Schwartz PJ, et al. (2020) Inherited cardiac arrhythmias. Nature reviews. Disease primers 6(1):58.

19. Greene HL (2000) Drug recalls underscore safety concerns. Health news (Waltham, Mass.) 6(5):4.

20. Roden DM (2004) Drug-induced prolongation of the QT interval. The New England journal of medicine 350(10):1013–1022.

21. Oshiro C, Thorn CF, Roden DM, Klein TE, & Altman RB (2010) KCNH2 pharmacogenomics summary. Pharmacogenetics and genomics 20(12):775–777.

22. Lu Z, et al. (2012) Suppression of phosphoinositide 3-kinase signaling and alteration of multiple ion currents in drug-induced long QT syndrome. Science translational medicine 4(131):131ra150.

23. Cohen IS, Lin RZ, & Ballou LM (2017) Acquired long QT syndrome and phosphoinositide 3-kinase. Trends in cardiovascular medicine 27(7):451–459.

24. Salata JJ, et al. (1998) A Novel Benzodiazepine that Activates Cardiac Slow Delayed Rectifier K^+^ Currents. Molecular pharmacology 54(1):220–230.

25. Liin SI, et al. (2015) Polyunsaturated fatty acid analogs act antiarrhythmically on the cardiac I_Ks_ channel. Proceedings of the National Academy of Sciences 112(18):5714–5719.

26. Matschke V, et al. (2016) The Natural Plant Product Rottlerin Activates Kv7.1/KCNE1 Channels. Cellular Physiology and Biochemistry 40(6):1549–1558.

27. Seebohm G, Pusch M, Chen J, & Sanguinetti MC (2003) Pharmacological activation of normal and arrhythmia-associated mutant KCNQ1 potassium channels. Circ Res 93(10):941–947.

28. Liin SI, Yazdi S, Ramentol R, Barro-Soria R, & Larsson HP (2018) Mechanisms Underlying the Dual Effect of Polyunsaturated Fatty Acid Analogs on Kv7.1. Cell Reports 24(11):2908–2918.

29. Boland LM & Drzewiecki MM (2008) Polyunsaturated fatty acid modulation of voltage-gated ion channels. Cell biochemistry and biophysics 52(2):59–84.

30. Goel A, et al. (2018) Fish, Fish Oils and Cardioprotection: Promise or Fish Tale? Int J Mol Sci 19(12).

31. Liu Y, et al. (2020) A PIP2 substitute mediates voltage sensor-pore coupling in KCNQ activation. Communications biology 3(1):385.

32. Yan C & Zou X (2016) MDock: An Ensemble Docking Suite for Molecular Docking, Scoring and In Silico Screening. Computer-Aided Drug Discovery, ed Zhang W (Springer New York, New York, NY), pp 153–166.

33. Huang SY & Zou X (2006) An iterative knowledge-based scoring function to predict protein-ligand interactions: I. Derivation of interaction potentials. Journal of computational chemistry 27(15):1866–1875.

34. Xu X, Ma Z, Duan R, & Zou X (2019) Predicting protein–ligand binding modes for CELPP and GC3: workflows and insight. Journal of Computer-Aided Molecular Design 33(3):367–374.

35. Hou P, et al. (2017) Inactivation of KCNQ1 potassium channels reveals dynamic coupling between voltage sensing and pore opening. Nature communications 8(1):1730.

36. Zaydman MA, et al. (2014) Domain-domain interactions determine the gating, permeation, pharmacology, and subunit modulation of the IKs ion channel. eLife 3:e03606.

37. Hou P, Shi J, White KM, Gao Y, & Cui J (2019) ML277 specifically enhances the fully activated open state of KCNQ1 by modulating VSD-pore coupling. eLife 8.

38. Lan X, et al. (2016) Grafting voltage and pharmacological sensitivity in potassium channels. Cell research 26(8):935–945.

39. Kim Y, Bang H, & Kim D (2000) TASK-3, a New Member of the Tandem Pore K+ Channel Family. Journal of Biological Chemistry 275(13):9340–9347.

40. Wu D, Pan H, Delaloye K, & Cui J (2010) KCNE1 remodels the voltage sensor of Kv7.1 to modulate channel function. Biophys J 99(11):3599–3608.

41. Antzelevitch C & Burashnikov A (2011) Overview of Basic Mechanisms of Cardiac Arrhythmia. Cardiac electrophysiology clinics 3(1):23–45.

42. Zimetbaum P (2012) Antiarrhythmic drug therapy for atrial fibrillation. Circulation 125(2):381–389.

43. Wang HS, et al. (1998) KCNQ2 and KCNQ3 potassium channel subunits: molecular correlates of the M-channel. Science 282(5395):1890–1893.

44. Lacerda AE, Kramer J, Shen KZ, Thomas D, & Brown AM (2001) Comparison of block among cloned cardiac potassium channels by non-antiarrhythmic drugs. European Heart Journal Supplements 3(suppl_K):K23–K30.

45. Morita H, Wu J, & Zipes DP (2008) The QT syndromes: long and short. Lancet 372(9640):750–763.

46. Chen YH, et al. (2003) KCNQ1 gain-of-function mutation in familial atrial fibrillation. Science 299(5604):251–254.

47. Estrada JC & Darbar D (2008) Clinical use of and future perspectives on antiarrhythmic drugs. Eur J Clin Pharmacol 64(12):1139–1146.

48. Hume JR & Grant AO (2015) Agents Used in Cardiac Arrhythmias. Basic & Clinical Pharmacology, 13e, eds Katzung BG & Trevor AJ (McGraw-Hill Medical, New York, NY).

49. Jost N, et al. (2005) Restricting excessive cardiac action potential and QT prolongation: a vital role for IKs in human ventricular muscle. Circulation 112(10):1392–1399.

50. Chen L, Sampson KJ, & Kass RS (2016) Cardiac Delayed Rectifier Potassium Channels in Health and Disease. Cardiac electrophysiology clinics 8(2):307–322.

51. Wang Q, et al. (1996) Positional cloning of a novel potassium channel gene: KVLQT1 mutations cause cardiac arrhythmias. Nat Genet 12(1):17–23.

52. Bellocq C, et al. (2004) Mutation in the KCNQ1 gene leading to the short QT-interval syndrome. Circulation 109(20):2394–2397.

53. Zipes DP (2015) Antiarrhythmic therapy in 2014: Contemporary approaches to treating arrhythmias. Nature reviews. Cardiology 12(2):68–69.

54. Frommeyer G & Eckardt L (2016) Drug-induced proarrhythmia: risk factors and electrophysiological mechanisms. Nature reviews. Cardiology 13(1):36–47.

55. Webb B & Sali A (2016) Comparative Protein Structure Modeling Using MODELLER. Current Protocols in Protein Science, (John Wiley & Sons, Inc.).

