## Supplementary Material for "Modulating the voltage sensor of a cardiac potassium channel shows antiarrhythmic effects"

### Supplementary Materials

### Materials and Methods

***In silico* compound screening.** At the time when we performed *in silico* screening, the structure of human KCNQ1 (hKCNQ1) was not yet solved. We therefore built the hKCNQ1 structure model based on the crystal structure of rat Kv1.2-Kv2.1 chimera (PDB entry: 2r9r) (56) using the program MODELLER (55). The model possesses a binding pocket near the region of the extracellular loop between S1 and S2 and the extracellular loop between S3 and S4 (Fig 1a). Using this hKCNQ1 model structure and our MDock docking software(33, 57, 58), we screened the compound library, Available Chemical Database (ACD, Molecular Design Ltd.), which consists of over 200,000 organic compounds.

After the structures of the hKCNQ1 channel were solved using cryo-EM (PDB entries: 6uzz, 6v00, and 6v01)(8), we used the $Ca^{2+}$-bound cryo-EM structure (6uzz) for the docking prediction of C28 interacting with hKCNQ1. The backbone root-mean-square deviation (RMSD) between our modeled VSD structure based on PDB entry 2r9r and the one in the cryo-EM structure (PDB entry: 6uzz, resolution 3.1 Å) is 4.4 Å. The overall structures of the two VSD models are very similar. However, the structure of the loop between S3 and S4 on VSD was missing in these cryo-EM structures. Therefore, the program MODELLER was employed to build the S3-S4 loop for the three cryo-EM hKCNQ1 structures. A total of 10 modeled structures were generated and used to study the detailed binding mode of C28 in hKCNQ1. C28 was docked to the modeled hKCNQ1 structures using the ensemble docking strategy implemented in our MDock docking program (33, 34). The predicted binding mode of C28 on VSD is shown in Fig 1A, 4E and Fig S4A, plotted with Chimera (59).

**Mutagenesis and expression of ion channels in *Xenopus* oocytes.** All mutations were made using overlap-extension PCR (polymerase chain reaction) with Pfu polymerase (Stratagene, CA). All PCR-amplified regions were confirmed by sequencing. The following ion channels and auxiliary subunits were studied: human KCNQ1 (UniProtKB/Swiss - Prot accession no. P51787), human KCNE1 (P15382), human Kir1.1 (P48048), hERG (Q12809), rat $K_V4.2$ (Q63881), human $Na_V1.5$ (Q14524), human $Na^+$ channel subunit β1 (Q07699), human $Ca_V1.2$ (Q13936), human $Ca^{2+}$ channel subunit β1 (Q02641), human $Ca^{2+}$ channel subunit $\alpha2/\delta1$ (P54289), and drosophila Shaker (P08510). The mMessage T3, T7 or SP6 polymerase kit (Invitrogen, Thermo Fisher Scientific, MO) was used to synthesize mRNA. 4-10 ng of mRNA encoding for the ion channel of interest in 46 nL volume was injected into *Xenopus laevis* oocytes (stage IV-V). For KCNQ1+KCNE1, mRNA mix at a weight ratio of 4:1 (KCNQ1:KCNE1) was injected. For $Na^+$ channels a 2:1 molar ratio (β1:$Na_V1.5$) and for $Ca^{2+}$ channels a weight ratio 1:1:1 (β1:$\alpha2/\delta1$:$Ca_V1.2$) was injected, respectively. Oocytes were then incubated in ND96 solution containing (in mM): 96 NaCl, 2 KCl, 1.8 $CaCl_2$, 1 $MgCl_2$, 5 HEPES, pH 7.6 supplemented with 2.5 mM Na - pyruvate and 1% penicillin - streptomycin, at 18°C for 3-7 days before recordings.

We also used different KCNQ1:KCNE1 ratios in our study. The effects of C28 on maximal conductance showed some quantitative differences but at all ratios the C28 effects on IKs did not show consistent differences.

All animal-related experimental protocols were approved by the Wahsington University in St. Louis Institutional Animal Care and Use Committee.

**Two-electrode voltage clamp recordings.** Microelectrodes were made of glass pipettes (Sutter Instrument, Item #: B150–117–10) using a puller (Sutter Instrument, P-97). The tip resistances were 0.5–3 MΩ when filled with 3 M KCl solution. Two-electrode voltage clamp (CA-1B amplifier, Dagan, Minneapolis, MN or GeneClamp 500B amplifier, Axon Instruments, CA) was used to record whole-oocyte ionic currents. The currents were sampled at 1 kHz and low-pass-filtered at 2 kHz. Patchmaster software (HEKA, Germany) was used to acquire data. The currents of Kir1.1 channels were recorded with a high $K^+$ bath solution containing (mM) 96 KCl, 4 NaCl, 1 $CaCl_2$, 2 $MgCl_2$, 10 HEPES, pH 7.3-7.4, The bath solution was ND96 solution for all other $K^+$ channels. The currents of $Ca_V1.2$ channels were recorded in 40 mM $Ba^{2+}$ bath solution containing (in mM): 40 $Ba(OH)_2$, 50 TEA-OH, 2 KOH, 5 HEPES, adjusted to pH 7.4 with methanesulphonic acid. C28 (alizarin blue black bg, MP Biomedicals, OH) and XE991 (10,10-bis(4-pyridinylmethyl)-9(10H)-anthracenone, Sigma-Aldrich, MO) were dissolved in bath solutions to the final concentration. Electrophysiological recordings were performed after ionic currents had become stable. All other chemicals were from Sigma-Aldrich. For measuring I-V relations of Kir1.1 the test voltage was -80 to 100 mV for 1s with increments of 10 mV. The test pulses were stepped from a holding potential of 0 mV and then stepped back to 0 mV. For all other $K^+$ channels and the $Ca^{2+}$ channel the holding potential was -80 mV, the test pulse was applied with 10 mV increments, which were followed with a repolarization pulse at -40 mV before returning to the holding potential. The durations of test pulses and the -

40 mV repolarization pulse vary for different channels to allow the activation and deactivation of the channel to reach steady states.

All electrophysiological recordings on oocytes and isolated ventricular and atrial myocytes were performed at room temperature (20-23 $^oC$).

**Cut-open oocyte clamp recordings.** Currents of $Na_V1.5$ channels were recorded using cut - open amplifier (CA - 1B; Dagan Corporation, MN). The internal solution was (mM): 105 NMG - Mes, 10 Na - Mes, 20 HEPES, and 2 EGTA, pH 7.4. The external solution contained (mM): 25 NMG - Mes, 90 Na - Mes, 20 HEPES, and 2 Ca - $Mes_2$, pH 7.4. The membrane potential was held at -100 mV. Currents were elicited by depolarization pulses in 10 mV increments with a 100 - ms - long prepulse and 50 - ms - long postpulse to - 120 mV.

**Voltage clamp fluorometry.** Oocytes were incubated for 30 min on ice in 10 μM Alexa 488 C5-maleimide (Molecular Probes, OR) for labeling in high $K^+$ solution (in mM) 98 KCl, 1.8 $CaCl_2$, 5 HEPES, pH 7.6. The labeling solution was then removed by washing oocytes three times with ND96 solution. Two electrode voltage clamp (CA-1B amplifier) was used to record whole oocyte currents. ND96 solution was used in the bath. The fluorescence signals were collected using a Pin20A photodiode (OSI Optoelectronics, CA), filtered using a FITC filter cube (Leica, Germany, for Alexa 488), and then amplified using a patch clamp amplifier (EPC10; HEKA, Germany).

**Analyses of ion channel function.** Conductance-voltage (G-V) relationships were obtained by measuring instantaneous amplitudes of tail currents following test pulses to various voltages. The G-V was normalized to the maximum G value. Fluorescence-voltage (F-V) relationships were derived by measuring the ΔF/F value at the end of test pulses and then normalized by dividing the maximum ΔF/F value. We used the Boltzmann equation to fit F-V and G-V curves:

$$G/G_{Max} = 1/(1 + \exp(-ze_o(V - V_{1/2})/kT)) = 1/(1 + \exp((V_{1/2} - V)/b)) \quad (1)$$

where $z$ represents the number of equivalent charges that move across the membrane electric field, $e_o$ is the elementary charge, $V$ is membrane potential, $V_{1/2}$ represents the voltage where $G/G_{Max}$ or $\Delta F/F_{Max}$ reaches 0.5, $k$ is Boltzmann's constant, $T$ is absolute temperature, and $b$ is the slope factor.

Dependence of G-V shifts ($\Delta V_{1/2}$) on C28 concentration was fitted to the Hill equation:

$$\theta = a+(b-a)/(1+(c/X)^d) \quad (2)$$

where $\theta$ is the expected response at dosage $X$, $a$ is the response when dosage = 0, $b$ is the maximum response or the stabilized response for an infinite dosage, $c$ is the dosage at which the response is 50% of maximum, which is also known as $EC_{50}$, and $d$ is the Hill Coefficient. In all our fittings Hill Coefficient was close to 1, and subsequently set to 1 for the final fitting.

**Computer models of canine ventricular action potentials.** The Hund-Rudy (HR) model of the canine membrane (non-propagating) ventricular action potential(60) is described by 30 ordinary differential equations (ODEs): 1 equation describing Vm, 6 equations describing sarcoplasmic and sarcoplasmic reticulum ion concentrations and 23 equations describing channel gating. (variable initial conditions can be found online at rudylab.wustl.edu).

Two modifications of the original Hund-Rudy model (MHR) were introduced to create simulations that more closely reflected experimental values for the sodium current: 1) the midpoint of inactivation was shifted from −70.3 mV, to −74.0 mV, and 2) maximal sodium conductance was changed to 14.2 mS/µF from 8.25 mS/µF. These modifications yielded a maximal upstroke velocity (Vmax) of 289 V/sec in MHR, more typical of the ventricular action potential than the 193V/sec generated by the original HR model. All computations were executed using the MATLAB® mathematical programming environment (see www.mathworks.com). All details of the computing methods are provided in the "on-line supplement: Computer Modeling" of a previous publication(61).

The simulations presented here only involved modifications of $I_{Ks}$ and $I_{Kr}$. Reducing $I_{Kr}$ was a means of creating a long Q-T syndrome *in silico*. The negative shift in the voltage dependence of $I_{Ks}$ activation was used as a means of testing the hypothesis that enhanced activation of $I_{Ks}$ might counteract the reduction in $I_{Kr}$, shortening the prolonged long Q-T action potential, while having relatively small effects in control conditions.

**Cardiac myocytes dissociation.** Guinea pigs, weighing about 500 g, were sacrificed with an overdose of sodium pentobarbitone (1 mL of 390 mg/ml) by abdominal injection. The hearts were perfused using a Langendorf procedure with 0.4 mg/ml collagenase (Worthington Biochemical Co.) at 37 $^{o}$C. The ventricular or atrial tissue was triturated to dissociate cells. Cells were then stored in KB solution containing (mM): KCl, 83; $K_2HPO_4$,

30; $MgSO_4$, 5; Na-Pyruvic Acid, 5; β-OH-Butyric Acid, 5; Creatine, 5; Taurine, 20; Glucose, 10; EGTA, 0.5; HEPES, 5; $Na_2$-ATP, 5. pH was adjusted with KOH to 7.2 (62) .

Mixed-breed dogs of either sex more than 12 months old were purchased from Marshall Farms. Dogs were sacrificed by intravenous injection of sodium pentobarbitone 100mg/kg. Canine ventricular cells were isolated using a modified Langendorf procedure by perfusing a wedge of left ventricle through a coronary artery with 0.5 mg/mL of collagenase (Worthington type 2) and 0.08 mg/mL of protease (Sigma Type XIV) for 12 to 15 minutes followed by tissue mincing (63). Cells were then stored in the KB solution described above.

All animal-related experimental protocols were approved by the Stony Brook University Institutional Animal Care and Use Committee.

**$I_{Ks}$ measurements in isolated cardiac myocytes.** An Axopath-1D amplifier (Axon Instrument) and the whole cell patch clamp technique were employed to record cell membrane currents. The cells were perfused with the external Tyrode solution we employed for measuring $I_{Ks}$ containing (mM): NaCl, 137.7; KCl, 8, NaOH, 2.3; $MgCl_2$, 1; Glucose, 10; HEPES, 5; $CdCl_2$, 1. pH = 7.4. C28 with various concentrations was added to the solution. The patch clamp electrode solution contained (mM): KOH, 140; HEPES, 20; KCL, 2; EGTA, 5; Mg-ATP, 5. pH = 7.2 adjusted with $HMeSO_3$. The patch electrodes had resistance of 2 – 3 MΩ with the filling electrode solution before sealing. The cells were held at -40 mV before the $I_{Ks}$ measurement. In this experimental condition, the Na-current, Na/Ca exchange current, Ca-current, Na/K pump current were eliminated, and the $I_{Ks}$ current was defined as the Chromanol 293B (10 µM, Sigma-Aldrich, MO) sensitive current. To measure the currents, the membrane potential was stepped up for 5s with 10 mV increments until 60 mV. The holding potential was -20 mV. The interval between the start of each measurement was 20 s to ensure the complete recovery of $I_{Ks}$ from the activation at the previous voltage step.

**Statistical tests of data.** All error bars in figures, including supplemental figures, represent standard error of mean (SEM), n = 3-15 unless otherwise specified. Figure 6E, Tukey–Kramer ANOVA test, n= 5-44; Figure 6G, t-test, n = 9-12. Statistical tests for supplemental results are described in each figure.

**Data Availability.** The datasets generated during and/or analyzed during the current study are available from the corresponding authors on reasonable request.

**Reference:**


56. Long SB, Tao X, Campbell EB, & MacKinnon R (2007) Atomic structure of a voltage-dependent K+ channel in a lipid membrane-like environment. *Nature* 450(7168):376-382.

57. Grinter SZ, Yan C, Huang SY, Jiang L, & Zou X (2013) Automated large-scale file preparation, docking, and scoring: evaluation of ITScore and STScore using the 2012 Community Structure-Activity Resource benchmark. *J Chem Inf Model* 53(8):1905-1914.

58. Yan C, Grinter SZ, Merideth BR, Ma Z, & Zou X (2016) Iterative Knowledge-Based Scoring Functions Derived from Rigid and Flexible Decoy Structures: Evaluation with the 2013 and 2014 CSAR Benchmarks. *J Chem Inf Model* 56(6):1013-1021.

59. Pettersen EF*, et al.* (2004) UCSF Chimera--a visualization system for exploratory research and analysis. *Journal of computational chemistry* 25(13):1605-1612.

60. Hund TJ & Rudy Y (2004) Rate dependence and regulation of action potential and calcium transient in a canine cardiac ventricular cell model. *Circulation* 110(20):3168-3174.

61. Lau DH*, et al.* (2009) Epicardial border zone overexpression of skeletal muscle sodium channel SkM1 normalizes activation, preserves conduction, and suppresses ventricular arrhythmia: an in silico, in vivo, in vitro study. *Circulation* 119(1):19-27.

62. Gao J, Mathias RT, Cohen IS, & Baldo GJ (1992) Isoprenaline, Ca2+ and the Na(+)-K+ pump in guinea-pig ventricular myocytes. *J Physiol* 449:689-704.

63. Zygmunt AC (1994) Intracellular calcium activates a chloride current in canine ventricular myocytes. *Am J Physiol* 267(5 Pt 2):H1984-1995.

64. Wallace AC, Laskowski RA, & Thornton JM (1995) LIGPLOT: a program to generate schematic diagrams of protein-ligand interactions. *Protein Eng* 8(2):127-134.

**FIGURE S1:**

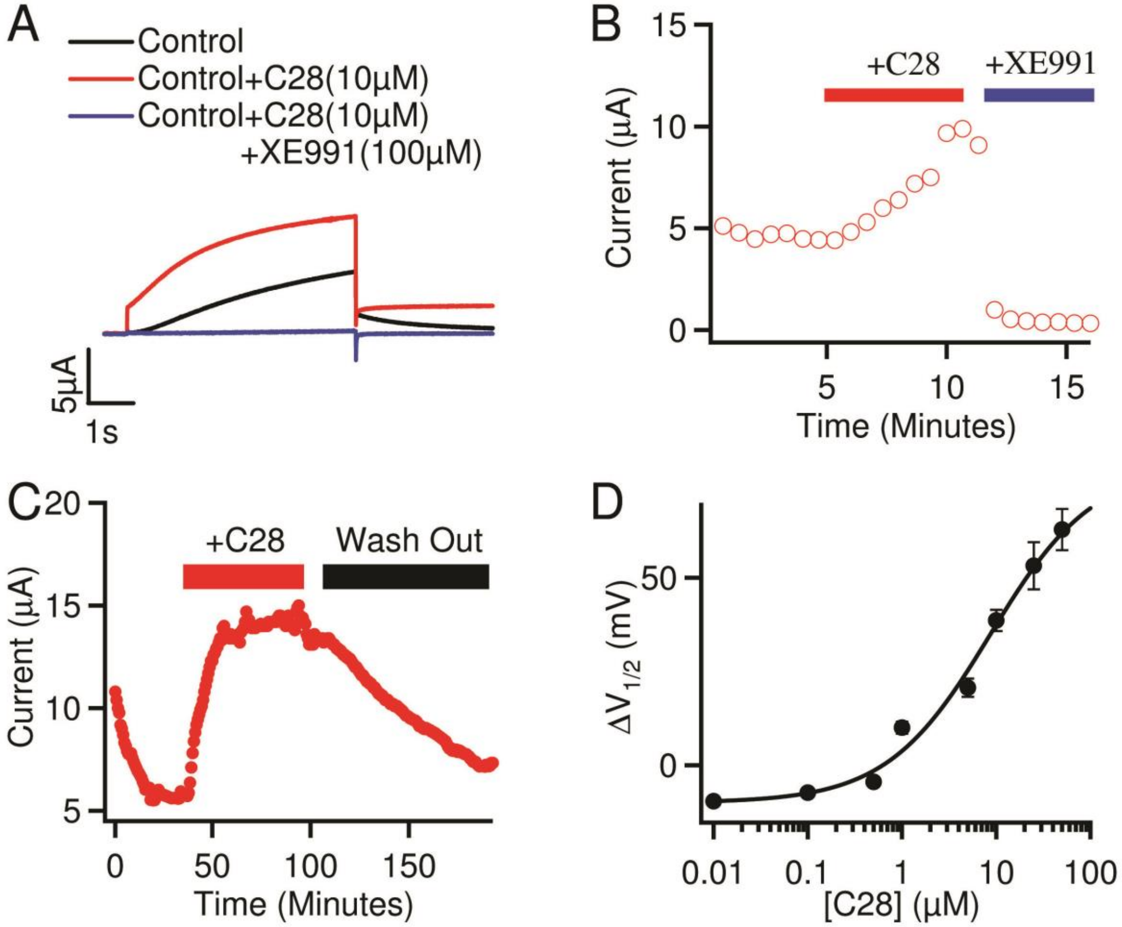
A
Control
Control+C28(10µM)
Control+C28(10µM)
+XE991(100µM)
5µA
1s
B
+C28
+XE991
Current (µA)
Time (Minutes)
C
+C28
Wash Out
Current (µA)
Time (Minutes)
D
ΔV1/2 (mV)
[C28] (µM)

**Figure S1. C28 effects on $I_{Ks}$ channels. A,** XE991, an inhibitor of KCNQ1 and $I_{Ks}$ channels, inhibits $I_{Ks}$ in the presence of C28. Voltage was stepped from a holding potential of -80mV to +40mV then to -40mV. **B,** $I_{Ks}$ in response to C28 (10 µM) and XE991 (100 µM) over time. **C,** $I_{Ks}$ in response to C28 (50 µM), and washing out of the C28 effect with ND96 solution. **D,** G-V shifts in response to application of C28 at various concentrations, which is the same data as in Fig 2D before correction. $\Delta V_{1/2} = V_{1/2}$ (control) - $V_{1/2}$ (in C28) < 0 in [C28] < 1 µM here because of fluctuations in $I_{Ks}$ during the long waiting time between the application of C28 and the start of G-V measurement. Please see B and C for examples of $I_{Ks}$ fluctuations and the long time needed for $I_{Ks}$ to stabilize after C28 applications. In Fig 2D the curve is moved up along the axis for $\Delta V_{1/2}$ such that $\Delta V_{1/2} = 0$ in 0.01 µM [C28]. This correction does not alter the fitting parameters for $EC_{50}$ or maximum $\Delta V_{1/2}$.

**FIGURE S2:**

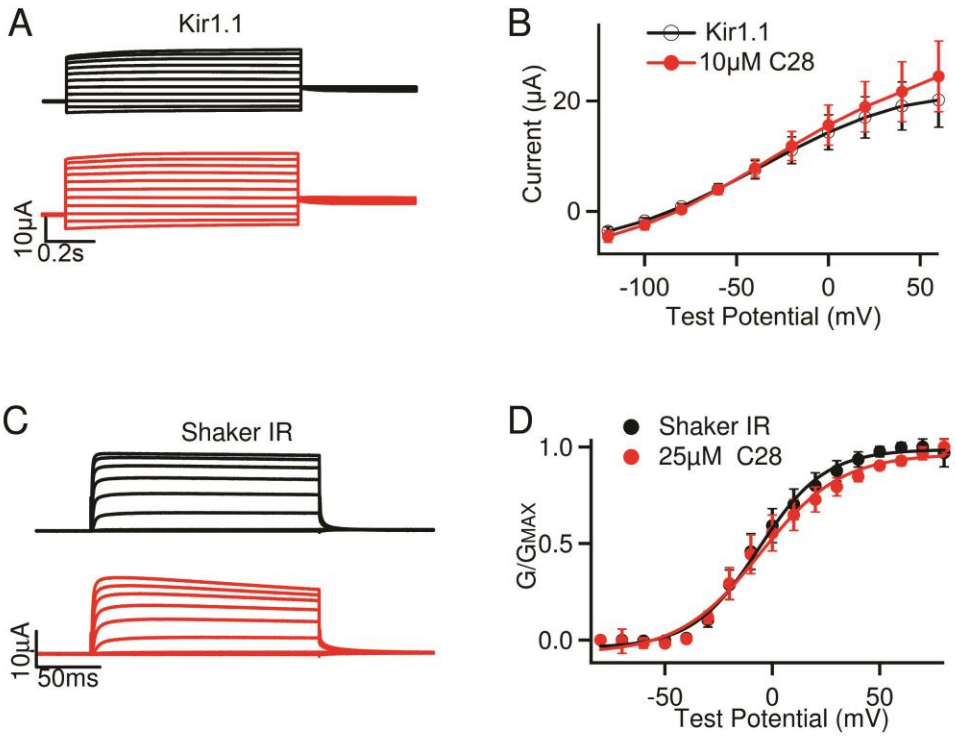

A
Kir1.1
10μA
0.2s
B
Kir1.1
10μM C28
Current (μA)
20
0
-100
-50
0
50
Test Potential (mV)
C
Shaker IR
10μA
50ms
D
Shaker IR
25μM C28
G/GMAX
1.0
0.5
0.0
-50
0
50
Test Potential (mV)

**Figure S2, C28 effects on channels that do not contain the KCNQ1 VSD. A**, Kir1.1 currents without (black) and with C28 (red) in response to voltages from -120 to 60 mV in 20 mV increments. Voltages before and after test pulses were -80 mV and -40 mV, respectively. **B,** Steady-state I-V relation with and without C28. **C,** Shaker-IR (IR - inactivation removed) currents without (black) and with 25 µM C28 (red). Test voltages were from -100 to 80 mV with 20 mV increments. Voltages before and after test pulses were -80 mV and -40 mV, respectively. **D,** G-V with and without C28.

**FIGURE S3:**

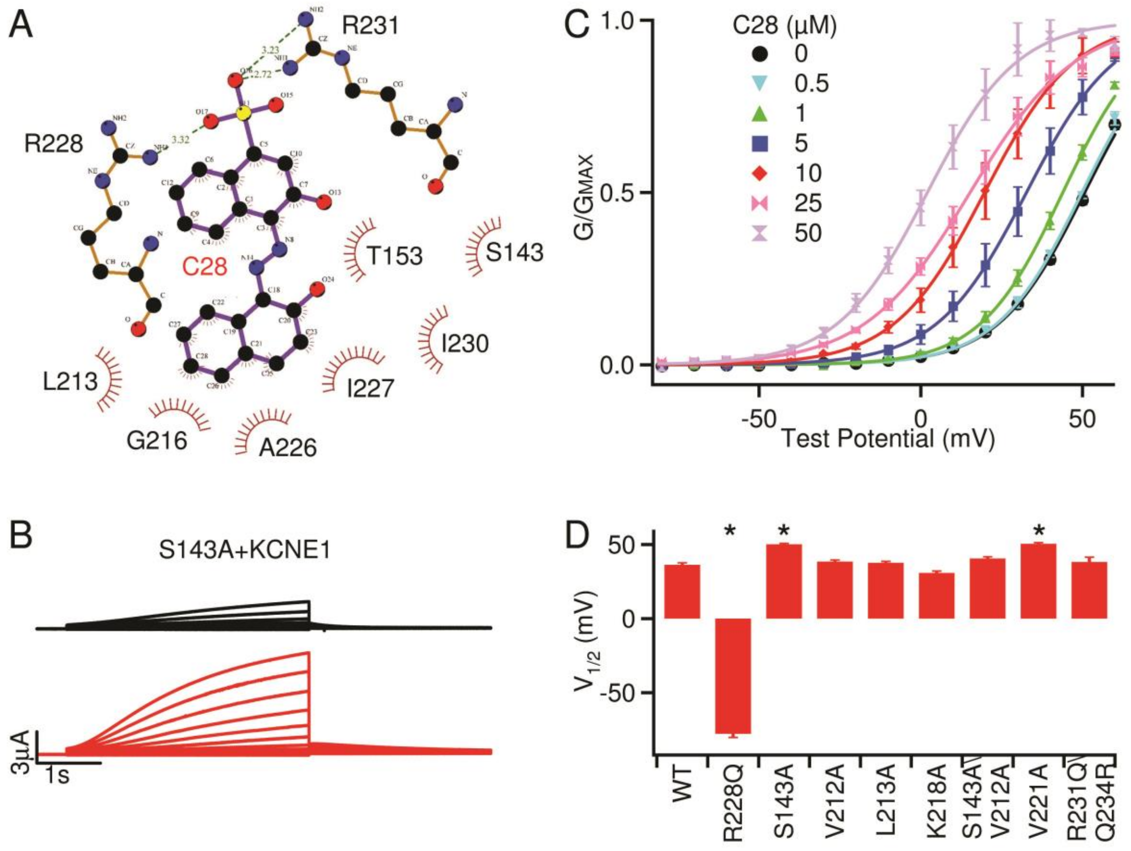
A
R231
R228
C28
T153
S143
I230
L213
I227
G216
A226
B
S143A+KCNE1
3µA
1s
C
C28 (µM)
0
0.5
1
5
10
25
50
G/GMAX
Test Potential (mV)
D
V1/2 (mV)
WT
R228Q
S143A
V212A
L213A
K218A
S143A\ V212A
V221A
R231Q\ Q234R

**Figure S3, Residues interacting with C28 predicted by docking and their role in $I_{Ks}$ opening.** **A,** Interaction details of C28 with human KCNQ1 channel in docking. The interacting residues were determined using the LIGPLOT program(64) based on the predicted complex structure (also shown in Fig. 4E). Two residues, R228 and R231, form hydrogen bonds with the ligand. Other residues are involved in hydrophobic contacts with the ligand. Experimentally confirmed critical residues for the ligand binding are represented in Fig. 4F. **B,** Mutation S143A KCNQ1 + KCNE1 (black) and with C28 (10 µM, red) from -80mV to 60 mV. The potentials before and after test pulses were -80 and -40 mV, respectively. **C,** G-V relationship of S143A KCNQ1 + KCNE1 with different C28 concentrations. Solid lines are fits to the Boltzmann relation. For the G-V with 0 [C28], $V_{1/2}$ and slope factor are 50.6 ± 2.3 and 13.4 ± 2.5. The shift of G-V ($\Delta V_{1/2}$) with different [C28] is shown in Fig 4F. **D,** Bar graph of $V_{1/2}$ of all mutants in the absence of C28. * $P<0.05$ compared to the WT $I_{Ks}$, Tukey–Kramer ANOVA test (n=16-26).

**FIGURE S4:**

A

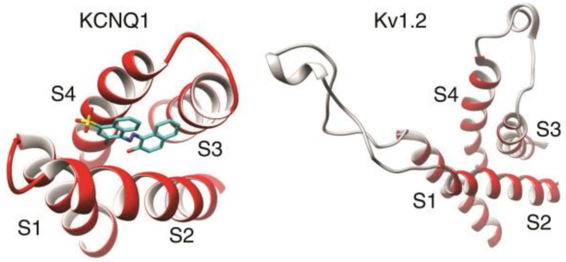


B

```
KCNQ1:          140 SVLST IEQYAAL ATGTL  197 PISIIDLIVVVASMVVLCVGS-- KGQVFATS
Herg:           420 YTAVF TP+24aa  PLAVV  496 GWFLIDMVAAIPFDLLIFGSG-- SEEL
Kv1.2:          178 VSFCL ET+37aa  FFIVE  255 -MNIIDIVAIIPYF-ITLGTELA EKPEDAQQ+6aa
Kv4.2:          200 IANVV ET+22aa  FFCLD  262 -MSIIDVVAILPYY-IGLVM--- TDNEDVS
Nav1.5 DI:      146 VFMAQ HDP+5aa  KYVEY  193 -WNWLDFSVIIMAYTTEFV---- DLGNVS
Nav1.5 DII:     732 LFMAL EHY+8aa  MLQVG  780 --NIFDSIIVILSLMELGLSRM- RMSNLS
Nav1.5 DIII:   1220 GALAF ED+11aa  LLEYA 1273 -WCWLDFLIVDVSLVSLVAN--- GFAE
Nav1.5 DIV:    1543 VTMMV ETD+8aa  ILAKI 1595 SWNIFDFVVVILSIVGTV----- QKYFFSPTLF
Cav1.2 DI:      139 VALAI YI+15aa  RVEYL  194 GWNLLDFIIVVVGLFSAIL---- EQATKADG+12aa
Cav1.2 DII:     539 LTIAS EH+13aa  ANKAL  587 LFNRFDCFVVCGGILETIL---- VETKIMSPLG
Cav1.2 DIII:    915 ISLAA ED+14aa  FDIVF  988 YFNILDLLVVSVSLISFGI---- QSSAINV
Cav1.2 DIV:    1254 ICLAM QH+13aa  LNMLF 1302 PWNVFDFLIVIGSIIDVIL---- SETNHYFC+44aa
                    S1    S1-S2 Loop S2        S3                      S3-S4 Loop
```

**Figure S4. Structural comparison among voltage gated ion channels. A,** Cryo-EM structure of KCNQ1 (PDB entry 6uzz) docked with C28 and crystal structure of Kv1.2. Note the different lengths of S1-S2 and S3-S4 loops. **B,** Sequence alignment of various ion channels. All 4 domains (DI-DIV) of Cav and Nav channels are included. The numbers indicate the position of the first amino acid in the channel sequence. The S1-S2 and S3-S4 loops are indicated by the number of amino acids (aa) that they contain if the length exceeds those in KCNQ1. Residues in red are those in KCNQ1 important for C28 interactions (Fig 4F) and the same corresponding residues in other channels. All sequences and topology of the channels were obtained from UniProt, then S3 segments were aligned using UniProt.

**FIGURE S5:**

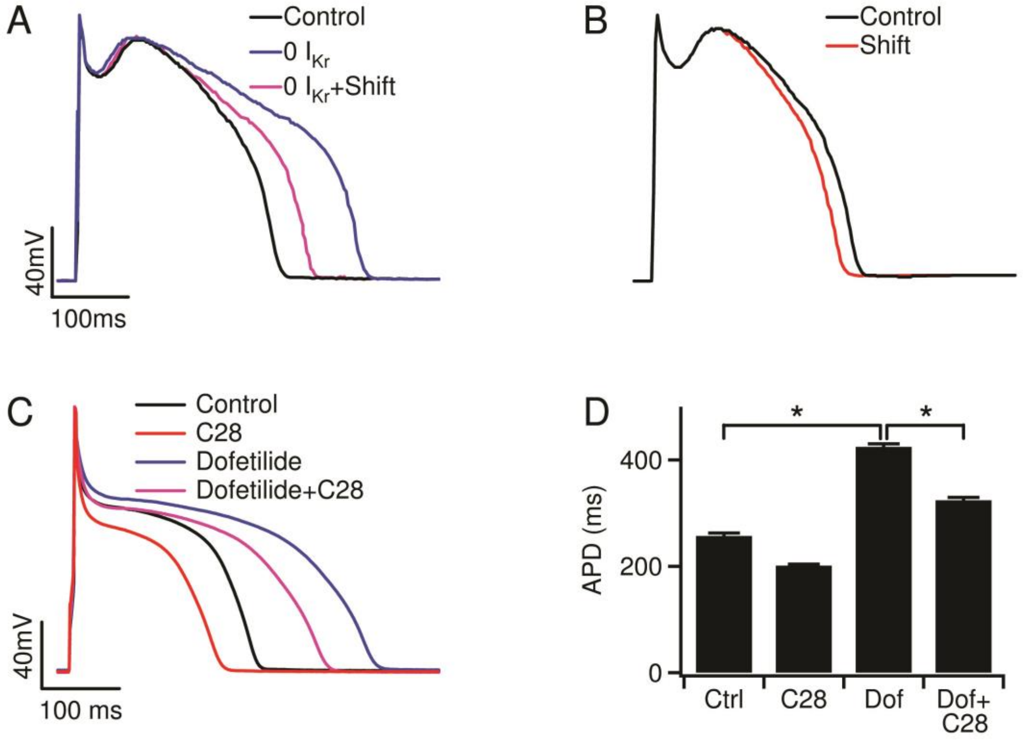
A
Control
0 I_Kr
0 I_Kr+Shift
40mV
100ms
B
Control
Shift
C
Control
C28
Dofetilide
Dofetilide+C28
40mV
100 ms
D
*
*
400
200
0
APD (ms)
Ctrl
C28
Dof
Dof+
C28

**Figure S5. An enhanced $I_{Ks}$ stabilizes action potentials in canine ventricular myocytes. A,B,** Simulation of the effects of a -20mV shift in the activation of $I_{Ks}$ on canine ventricular action potential model (60) with a total block of $I_{Kr}$ (**A**) or the control action potential (**B**). **C,** The effects of 10 µM C28 on the APD of canine ventricular myocyte in control and in the presence of 1 µM dofetilide, which blocks $I_{Kr}$. **D,** Average results for control and with 1 µM dofetilide. * P<0.05, student *t* test (n=7 or 8).

**FIGURE S6:**

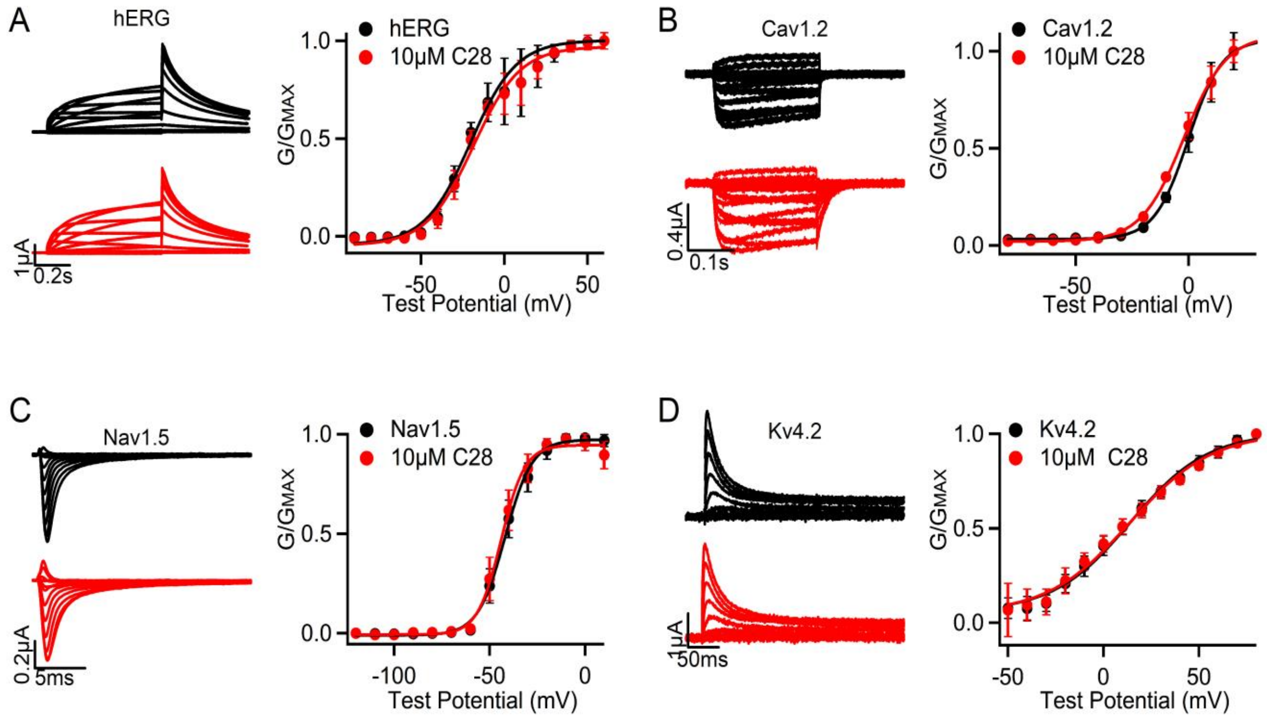
A
hERG
1μA
0.2s
hERG
10μM C28
G/GMAX
1.0
0.5
0.0
-50
0
50
Test Potential (mV)
B
Cav1.2
0.4μA
0.1s
Cav1.2
10μM C28
G/GMAX
1.0
0.5
0.0
-50
0
Test Potential (mV)
C
Nav1.5
0.2μA
5ms
Nav1.5
10μM C28
G/GMAX
1.0
0.5
0.0
-100
-50
0
Test Potential (mV)
D
Kv4.2
1μA
50ms
Kv4.2
10μM C28
G/GMAX
1.0
0.5
0.0
-50
0
50
Test Potential (mV)

**Figure S6. C28 effects on other currents activated in cardiac action potentials.** Current traces for indicated channels without (black) and with (red) 10 μM C28 (left) and G-V relations without and with C28 (right). **A**, hERG, voltages from -90 to 30 mV with 10 mV increments. $\Delta V_{1/2}$ of G-V relations ($V_{1/2}$ of C28 - $V_{1/2}$ of control): 1.7 ± 2.5 mV, n = 3, p = 0.82, Student T test. **B**, Cav1.2, voltages from -80 to 20 mV with 10 mV increments. $\Delta V_{1/2}$: -2.0 ± 0.5 mV, n = 4, p = 0.51. **C**, Nav1.5, voltage from -120 to 20 mV at 10 mV increments. $\Delta V_{1/2}$: -2.0 ± 0.9 mV, n = 4, p = 0.77. **D**, Kv4.2, voltages from -50 to 80 mV in 20 mV increments. $\Delta V_{1/2}$: -1.1 ± 1.1 mV, n = 3, p = 0.65.

**FIGURE S7:**

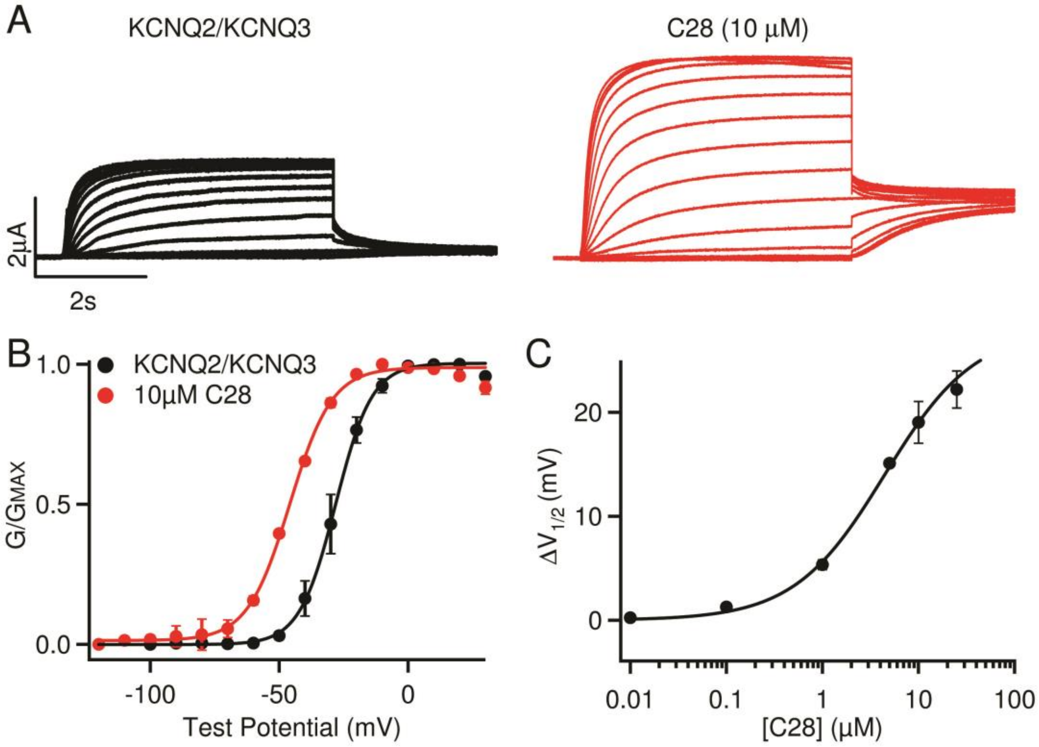

A
KCNQ2/KCNQ3
C28 (10 μM)
2μA
2s
B
KCNQ2/KCNQ3
10μM C28
1.0
0.5
0.0
G/GMAX
-100
-50
0
Test Potential (mV)
C
20
10
0
ΔV1/2 (mV)
0.01
0.1
1
10
100
[C28] (μM)

**Figure S7, C28 effects on KCNQ2/KCNQ3. A**, KCNQ2/KCNQ3 currents without (black) and with C28 (red) in response to voltages from -120 to 20 mV in 10 mV increments. Voltages before and after test pulses were -80 mV and -40 mV, respectively. **B,** G-V relation with and without C28. Solid lines are fits to the Boltzmann relation with $V_{1/2}$ and slope factor (mV) for control: -26.4 ± 2.4 mV and 7.1 ± 0.5 mV; and for 10 µM C28: -45.9 ± 2.4 mV and 8.2 ± 0.3 mV. **C,** The change in $V_{1/2}$ of G-V relations depends on C28 concentration, with an $EC_{50}$ of 4.3 µM and a maximal shift 27.5 mV.

**FIGURE S8:**

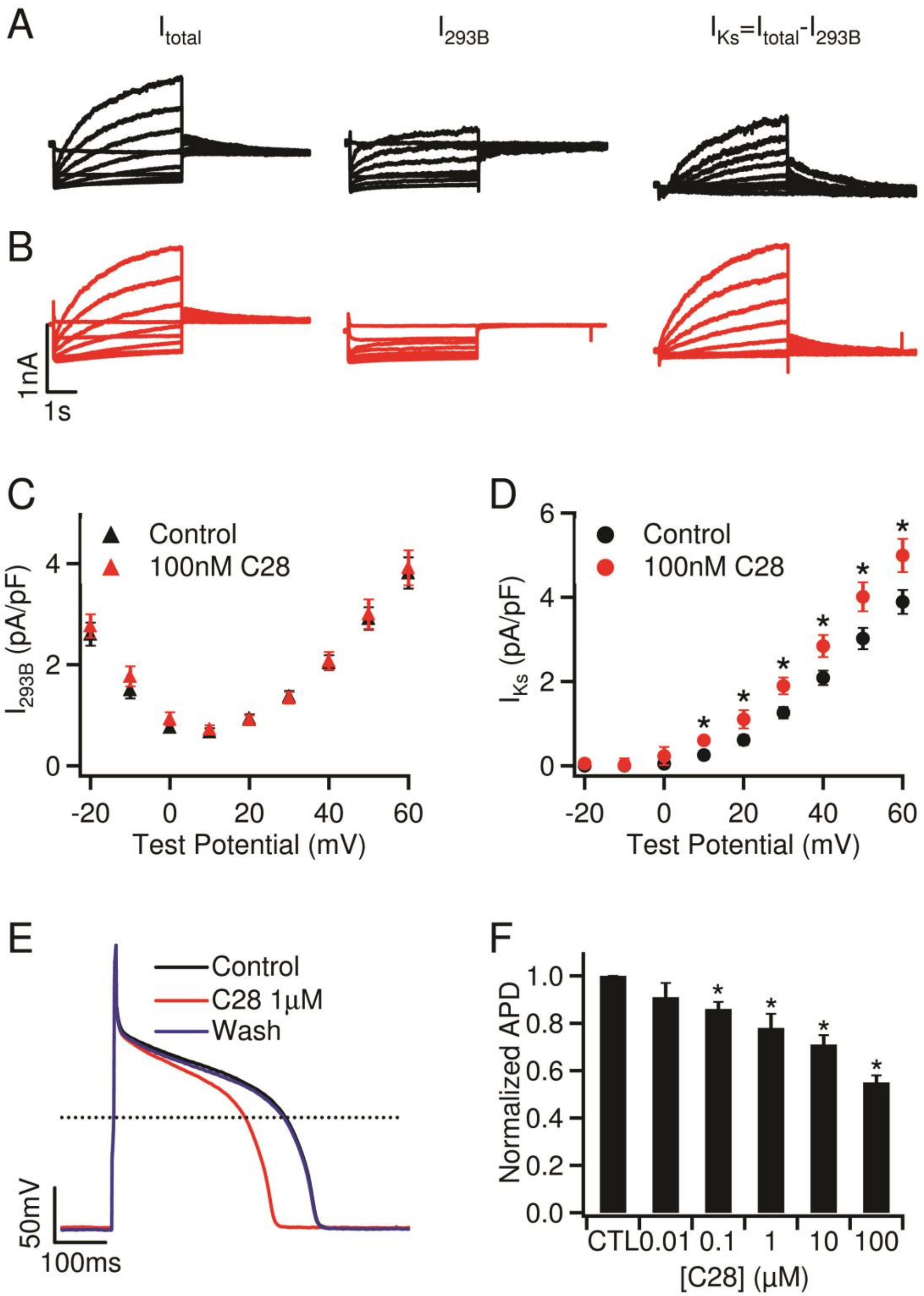
A
$I_{total}$
$I_{293B}$
$I_{Ks}=I_{total}-I_{293B}$
B
1nA
1s
C
Control
100nM C28
$I_{293B}$ (pA/pF)
Test Potential (mV)
D
Control
100nM C28
$I_{Ks}$ (pA/pF)
Test Potential (mV)
E
Control
C28 1μM
Wash
50mV
100ms
F
Normalized APD
CTL 0.01 0.1 1 10 100
[C28] (μM)

**Figure S8. C28 effects on $I_{Ks}$ and action potentials in GP ventricular myocytes. A, B,** Current traces recorded from GP ventricular myocytes, control (**A**) and with 100 nM C28 (**B**). Currents were recorded at various voltages (see **C**) without ($I_{total}$) and in the presence of 10 µM Chromanol 293B ($I_{293B}$), which is an inhibitor of $I_{Ks}$. The subtraction of $I_{293B}$ from $I_{total}$ gave rise to $I_{Ks}$. **C,** Current-voltage relations in the presence of 10 µM Chromanol 293B. **D,** Current-voltage relations of $I_{Ks}$. * P<0.05 compared to control, Student T-test. **E,** Sample action potential recordings. The current clamp stimulus was 180 pA in amplitude, 10 ms in duration at 1Hz frequency. **F,** The average APD in control and in a range of C28 concentrations. * P<0.05 compared to control, Tukey–Kramer ANOVA test (n=8-57).
